# Correlated Drug Action as a Baseline Model for Combination Therapy in Patient Cohorts and Cell Cultures

**DOI:** 10.1101/2021.05.23.445343

**Authors:** Adith S. Arun, Sung-Cheol Kim, Mehmet Eren Ahsen, Gustavo Stolovitzky

## Abstract

Identifying and characterizing the effect of combination therapies is of paramount importance in various diseases, including cancer. Various competing null models have been proposed to serve as baselines against which to compare the effect of drug combinations. In this work, we introduce Correlated Drug Action (CDA), a baseline model for the study of drug combinations in both cell cultures and in patient populations. CDA assumes that the efficacy of pairs of drugs to be used in a combination may be correlated, that is, if the efficacy of a drug in a given patient or cell is high (low), then the efficacy of the other drug may also be high (low) in the same patient or cell. Our model can be used in the temporal domain (temporal CDA or tCDA) to explain survival curves in patient populations, and in the dose domain (dose CDA or dCDA), to explain dose-response curves in cell cultures. At the level of clinical trials, we demonstrate tCDA’s utility in identifying possibly synergistic combinations and cases where the combination can be explained in terms of the monotherapies. At the level of cells in culture, dCDA generalizes null models such as Bliss independence, the Highest Single Agent model, the dose equivalence principle, and is consistent with what should be expected in sham combinations. We demonstrate the applicability of dCDA in assessing combinations in experimental MCF7 cell-line data by introducing a new metric, the Excess over CDA (EOCDA).

## Introduction

Combination therapy, the principle of treating a patient with multiple drugs either simultaneously or in succession, is a widely used strategy for treating conditions ranging from HIV/AIDS to cancer [1–4]. In the case of cancer, Frei and Freirich reasoned that cancer cells could mutate and become resistant to one chemotherapy but would be less likely to become resistant to multiple different anti-cancer chemotherapy agents [5]. Since that pioneering work, combination therapies have become the *de facto* approach for treating cancer. However, quantifying the effects of combination therapies can be experimentally inefficient given the number of possible combinations and resources such as participants, physicians, and assays needed. Thus, it is important to develop models that can characterize the effects of drugs acting in combination from the effects of the drugs acting independently. Null models, that is, models that assume the combination effect results from some sort of superposition of the individual effects of the monotherapies, can serve as a baseline for the expected effects of drugs applied in combination [6]. For example, such models can be used as a reference to determine if a combination is more or less effective than expected.

The effect of combinations has been studied at two fundamentally different levels: cultured cells (or *in-vitro*) and living organisms (or *in-vivo*). Cell lines and organoids fit into the former category whereas Patient Derived Xenografts (PDXs) and human clinical trials are examples of the latter. At the cell culture level, research typically focuses on the dose response at a fixed time point after drug administration; at the living-organism level, research tends to focus on survival time at fixed doses. We will refer to models at the level of cultured cells as dose-space models since the primary variable to change is the dose of the monotherapies in the combination. Similarly, models describing patient populations will be referred to as temporal models given that the main endpoint is survival times.

Recently, Palmer and Sorger [7] described effects of drug combinations in survival analysis in patient cohorts based on the principle of Independent Drug Action (IDA), which was first proposed by Bliss [8] with the name of Independent Joint Action in the context of the toxicity of poisons acting in combination. In its most basic form, IDA postulates that even when two drugs are given in combination, they act independently, each drug acting as if the other drug hadn’t been administered. Independent joint action, however, doesn’t imply statistical independence. As clearly stated by Bliss in [8], the susceptibility of the target (patient, cell or animal) to one component of a combination may or may not be correlated to the susceptibility to the other component. However Bliss didn’t provide any detail on how to compute this association. In [7], the authors used the Spearman correlation between the survival times that patients would have had if they had been treated with the individual drugs to quantify this association, and posited that the survival of a patient treated with a combination of drugs is equal to the survival that the patient would have had if they had been treated with the monotherapy with longer survival time. This independence-based baseline model can be useful to model combinations in patients under the assumption that the mode of action of one of the drugs does not affect the mode of action of the other drug. [7] studied the role of correlation between individual drug responses and concluded that benefit of a combination is the highest when drug responses are uncorrelated and no benefit is expected above the better of the two drugs when the responses are perfectly correlated.

In the current work we build on [8] and derive a closed mathematical formula for predicting the distribution of survival times for a combination in terms of the distribution of survival times for the monotherapies, under the assumption of independent drug action. Our formula is an approximation that interpolates between three special cases of the correlation between the survival times of a patient to the two drugs in the combination: when the correlation is -1, 0, 1. In these special cases it is possible to write the mathematical expression of the distribution of survival times for the combination. Note that unlike earlier work, our formula also allows for a negative association between individual drug responses. To the best of our knowledge, negative associations between individual drug responses have not been studied before. However, our analysis of IDA in the dose space suggests that negative correlations might exist. The joint distribution of survival times is in principle unknown. Interestingly, we prove in the Supplement that for a special case of joint probability distribution of survival times, our closed-form approximate formula turns out to be exact if the correlation is the Spearman correlation (see *Integrated expression for the survival probability under the tCDA model* in Supplement). We show in the supplement that the distribution of survival times under a combination depends very mildly on the joint distribution of survival times, and exact formula is a good approximation for a wide range of joint probability distributions of survival times.

Because of the essential role played by the correlation between individual drug responses as discussed above, we think that an appropriate name for this modeling framework should be Correlated Drug Action (CDA) to emphasize the correlation between susceptibilities to monotherapies in a combination. It should be emphasized that the drugs still are assumed to act independently; however we feel that calling the framework independent drug action may mask the fundamental role played by the correlation between responses to each drug. In the context of studying the survival times in patient populations, we will call our framework as the temporal CDA or tCDA. Our work differs from that of [7] in that, rather than using heavy simulation to fit the correlation between susceptibilities between drugs, we use our closed-form solution which allows for a computationally faster and easily scalable model estimation and downstream statistical testing, which we describe in the Supplement. We use this statistical framework to test for responses to combination that are statistically significantly different from the CDA mechanism such as synergistic and antagonistic drug interactions.

Besides recognizing independent joint action as a potential mechanism of drug action in a combination, Bliss [8], also proposed a second mechanism he called Similar Joint Action (SJA). In SJA, the drugs produce similar but independent effects so that one drug can be compensated by a constant proportion for the other, and variations in individual susceptibility to the two drugs are completely correlated. Since [8], various models have been proposed to rationalize the joint action of two drugs in dose space. The two most popular are the Bliss independence [9] model and the Loewe additivity model [10]. The Bliss independence model assumes IDA with no correlation between the responses to the two drugs, whereas the Lowe additivity model assumes the SJA. Since then various experimental papers have used these these two models to assess the synergy and antagonism of combination therapies without much emphasis on the fundamentals used to derive those formulas. Over time these two reference models have become competitors causing a big confusion in the field. Therefore, extrapolating the temporal ideas about correlations between survival times of each patient in a cohort under each monotherapy to the dose-space, we propose the dose CDA or dCDA model that describes the effect of combinations in cell cultures in terms of the dosages of each monotherapy that kill each cell in culture after a given treatment time. From simple assumptions, we derive a closed-form approximate expression for the viability of a cell culture to a combination. This expression interpolates between the Bliss independence model, valid in the case of no association between the susceptibilities of the culture to the drugs, and a sham-compliant Highest Single Agent (HSA) [11] model (a generalization of the Loewe additivity model) valid in the case of complete association of susceptibilities. Similar to tCDA, in dCDA, we assume a generalized, a priori unknown, joint distribution between the lethal doses of a cell to each drug in a combination, and show that for a specific joint distribution the closed-form expression is exact and the association reduces to the Spearman correlation between lethal doses. The range of correlation values ([−1, 1]) allows for a gamut of possible outcomes for the effect of a combination under dCDA ranging from the Bliss independence model (when the Spearman correlation is 0) to a sham-combination-compliant HSA (when the Spearman correlation is 1).

We must make clear that the correlation described in either model cannot be directly measured experimentally because it is not possible to treat a given cell with one monotherapy and then reset to the pre-initial monotherapy conditions and administer the other monotherapy to the same cell. Thus, the correlation is a latent variable that must be estimated from the data. Our modeling framework is flexible and can be easily extended to multi-drug therapies as long as we have sufficient data.

We apply the tCDA model to public oncology clinical trial data in order to demonstrate its ability to explain the effect of clinical combination therapies and test for combinations that cannot be explained with the tCDA model. When the survival distribution of the combination can be explained by the tCDA model, the correlation parameter may have a biological interpretation. The correlation can potentially inform whether there may exist a biomarker or patient covariate that can decouple the patient population into sub-populations that can benefit more from one monotherapy over the other, or those that realize the benefit of the combination.

Rather than introducing an entirely new model in an area that has seen much debate [12], the dose-space CDA baseline model interpolates different individual concepts (Bliss independence, HSA, the dose equivalence principle and sham combination compliance) and creates a spectrum of from independent joint action to fully correlated joint action between these established models. This semi-unification occurs rather naturally from the basic premises of the approach used to develop the CDA model (See *Formulation of the Dose-Space Correlated Drug Action Model* in Supplement). We show that the dose-space CDA model is useful in assessing whether there are synergistic effects in a given combination tested on cultured cells. To do that, we introduce a new performance metric we call Excess over CDA (EOCDA) and associated statistical framework. Further, using experiments in a cell line (MCF7) and several combinations, we propose an approach for identifying doses at which the combinations produce outlier results that may be interpreted as synergistic combinations.

## Results

### Temporal Correlated Drug Action as a Model to Explain the Effect of Drug Combinations in Human Clinical Trials

To illustrate the use of our tCDA model, we next discuss its application to different combination therapy clinical trials. A detailed discussion of the simulation methods motivating the tCDA model, the closed-form expression underlying tCDA, and the statistical testing framework for assessing whether a given combination cannot be explained by the tCDA model can be found in the sections *Detailed examination of the tCDA model with Herceptin and chemotherapy combination trial data* and *Additional examples of tCDA model applied to combination trial data* of the Supplement. For the 18 different combinations tested in clinical trials and retrieved from the literature (See Methods and Supplementary Table S1), we estimated the optimal tCDA predicted survival curve and assessed whether the tCDA model adequately described the observed combination (see Fig. 1A). We developed a procedure for estimating the correlation between single drug responses (*ρ*) using our closed-form mathematical expression for the tCDA model (see *Formulation of the temporal correlated drug action model* in Supplement) and a framework to estimate the statistical significance of the model-to-data fit (see section *A statistical test to compare the analytical tCDA model to the measured survival probability*). These methods are quite fast and scalable. In our analysis, we adjust for testing multiple hypotheses by applying a Benjamaini-Hochberg correction to the individual significance level of 0.05. In 66% of the tested combinations (12 of the 18 trials), we failed to reject the tCDA model (p-value > 0.003) (Fig. 1A). In such cases, the estimated combination PFS curve closely tracks the experimentally observed combination (1B-F; Figs. S3D, S4C, S4D, S5A, S6A, S6C). We also used tCDA to analyze combinations composed of more than two drugs (e.g., Fig. 1F). In 33% of the tested combinations (6 of the 18), we reject the tCDA null model (Fig. 1A). These represent the combinations that perform differently than expected under the tCDA model. We observed that the tightness of the fit between the tCDA model and the actual combination can act as a red herring, distracting us from the fact that this combination is statistically significantly different than the one predicted by tCDA, as suggested by our statistical significance framework. This phenomenon can be noted in the Dabrafenib and Trametinib combination in metastatic BRAF-mutant cutanous melanoma (Fig. S6B). This encouragingly suggests a synergistic interaction which is supported by the mechanism of action of both drugs. Trametinib is a selective inhibitor of MEK1/2 activity and Dabrafenib is a potent inhibitor of BRAF and CRAF [13, 14]. Both BRAF and MEK are in the same pathway where BRAF is upstream of MEK, so drugs that inhibit these two targets provide a classical example of synergy. There are more obviously synergistic combinations, like the 5-FU and Oxaliplatin combination for advanced pancreatic cancers for which the statistical test (p-value *<* 10^−5^) supports the visual fit and mechanistic understanding of the combination (Fig. S4A).

**Figure 1:**
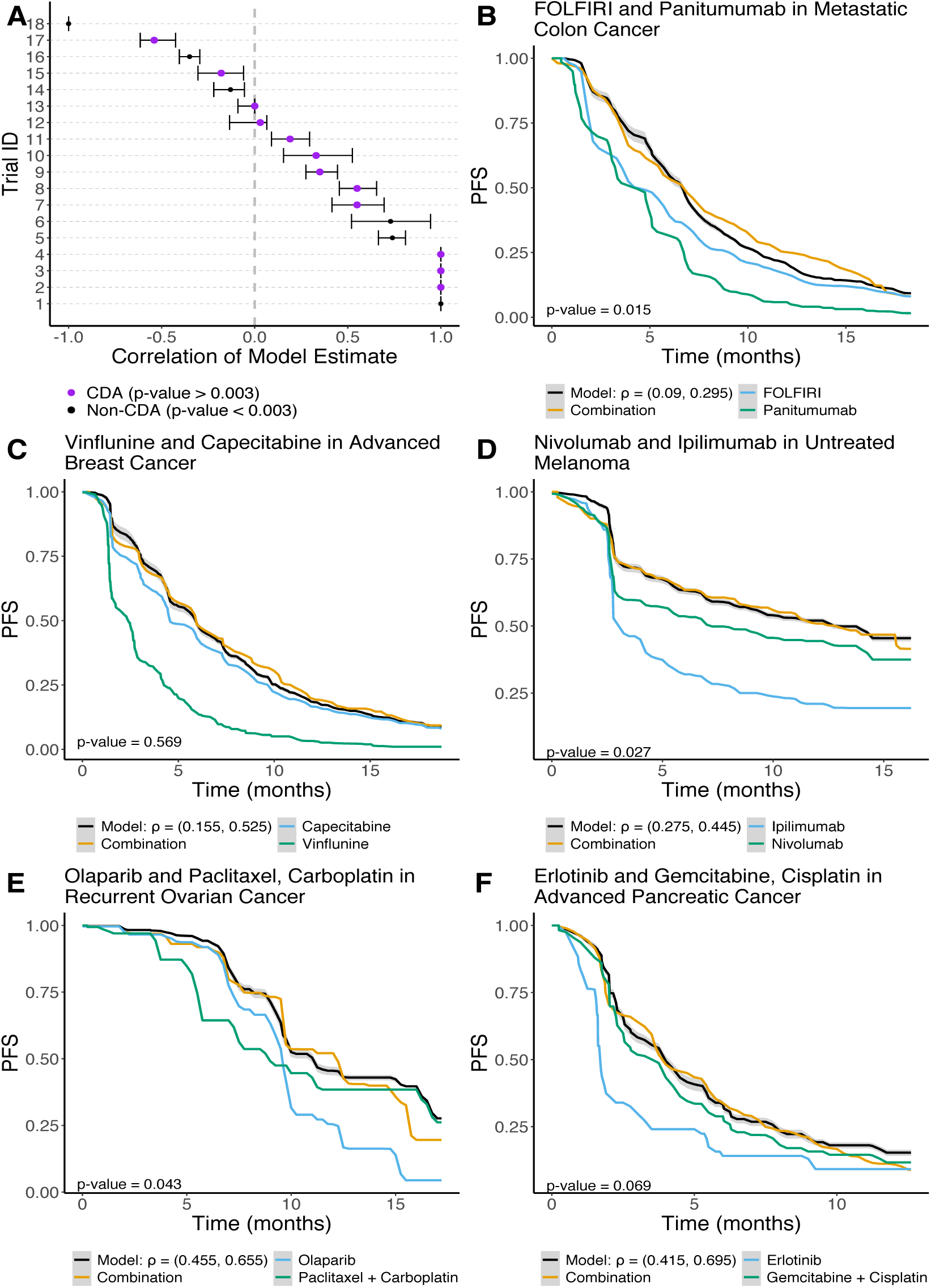
Temporal Correlated Drug Action (tCDA) model results. **A)** Optimal Spearman’s correlation and associated 95% confidence interval. Purple represents trials where the effect of the combination can be explained sufficiently well under the tCDA model. The trial ID’s reference the tested clinical trials, and a lookup table can be found in Supp. Table 1. **B)** tCDA model estimate for the combination of FOLFIRI and Panitumumab in metastatic colon cancer. **C)** tCDA model estimate for the combination of Vinflunine and Capecitabine in advanced breast cancer. **D)** tCDA model estimate for the combination of Nivolumab and Ipilumumab in previously untreated melanoma. **E)** tCDA model estimate for the combination of Olaparib and Paclitaxel, Carboplatin in recurrent ovarian cancer. **F)** tCDA model estimate for the combination of Erlotinib and Gemcitabine, Cisplatin in advanced pancreatic cancer.

### Dose Correlated Drug Action as a Model to Explain the Effect of Drug Combinations in Cell Cultures

In this section we extend the principle of independent drug action, originally presented to address survival of cancer patients treated with drug combinations, to the problem of determining cell viability in response to drug combinations in cell cultures and in dose space. To do so we apply the same principles that were applied for tCDA but rather than asking for the survival time of a patient under each of two drugs, we will consider the lethal dose of a cell to each of two drugs after a fixed treatment time has elapsed (typically 24, 48 or 72 hours). The lethal dose, which depends on the treatment time, is a property of each cell in response to a drug. Each cell has different lethal doses for different drugs. A cell treated with a given drug at a specified dose and for a specified treatment time will be found dead (resp. alive) after that time if its lethal dose is below (resp. above) the applied dose. The extension of tCDA to cell cultures treated with a pair of drugs postulates that a cell in the culture will be viable after the treatment time has elapsed if the lethal dose of that cell for each of the monotherapies in the combination is larger than the dose of corresponding drug in the pair (see *Integrated expression for the viability of cells under the dCDA model* in the Supplement). Therefore cells that had at least one of its lethal doses smaller than the applied doses in the combination will be found dead at the targeted treatment time. An important assumption in what follows is that the lethal doses of the two drugs in a combination may be correlated. The origin of this correlation could be some similarity in the mechanism of action and/or the action of some confounding variables. In cell cultures we can think that specific characteristics of a cell such as its number of mitochondria [15], its size, its protein content, etc., could be a confounder that simultaneously affect the lethal doses of each of the two drugs. The dCDA model can be expressed as the following closed-form expression (see *Dose-Space Correlated Drug Action* in Supplement)

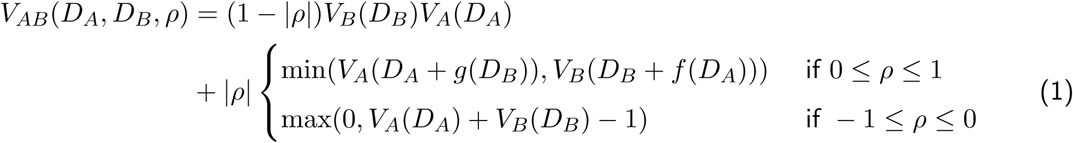

where *V*_*AB*_(*D*_*A*_, *D*_*B*_, *ρ*) is the dCDA predicted viability of the cell culture for the combination of drugs A and B at doses *D*_*A*_ and *D*_*B*_ when the Spearman’s rank correlation of the lethal doses to A and B is *ρ, V*_*A*_(*D*_*A*_) and *V*_*B*_(*D*_*B*_) are the dose response curves for the cell culture treated with drugs A and B independently. The function *f* (*D*_*A*_) (resp. *g*(*D*_*B*_)) is the equivalent dose of B (resp. A) that would produce the same effect in the culture viability as the dose *D*_*A*_ of A (resp., the dose *D*_*B*_ of B) did, and are discussed in the Supplement. For *ρ* = 0 the dCDA model reduces to the Bliss independence model. For *ρ* = 1 the dCDA model reduces to a modified version of the HSA model that is sham compliant which is a generalization of Loewe additivty. Indeed, when a drug is ‘combined’ with itself the lethal doses of a cell to the same drug would result in a Spearman’s correlation value of *ρ* = 1.

The dCDA model requires that we find the optimal parameter *ρ* across a range of different doses in order to estimate the combination viability in the dCDA framework (See *Fitting the dCDA model to the data* in Supplement). Across the 26 experimentally tested combinations in MCF7 cells discussed below (Supp. File 2), the average Spearman’s correlation estimate is 0.03 and ranges from -0.14 to 0.21 (Fig. 2A). For a given combination estimate, we assess whether the viability of a combination predicted by the dCDA model sufficiently describes the observed viability of the combination using a two-sample paired t-test (Figs. 2D, F; See *Fitting the dCDA model to the data* in Supplement). We chose this method to minimize false negative rates, even when it results in some extra false positive CDA abiding combinations, for which we should have rejected the CDA null hypothesis. To compensate for this increase in false positive combinations, we will introduce later on a “local analysis” that flags specific doses of combinations as not abiding by the dCDA model.

**Figure 2:**
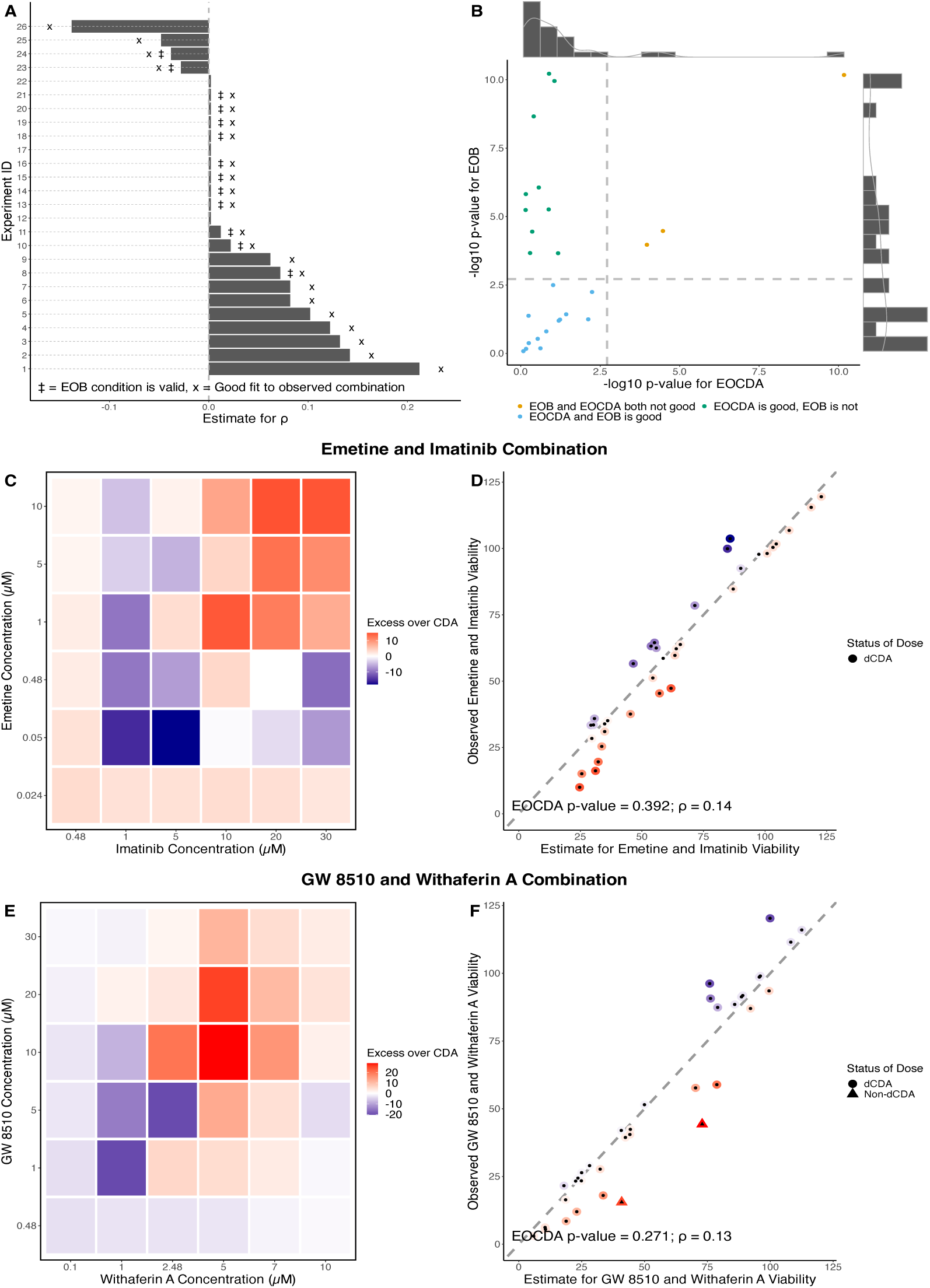
Dose-space Correlated Drug Action (dCDA) model results. **A)** Optimal Spearman’s correlation estimate for each *in vitro* drug combination tested on MCF7 cells under the dCDA model. Double-dagger indicates a good fit (p-value > 0.01), and star indicates that Excess Over Bliss (EOB) is a valid metric (p-value > 0.01). The trial ID’s correspond to a given drug combination (Supp. Table 2). **B)** Joint distribution of p-value statistics for Goodness of Fit and EOB conditions (Dashed lines = p-value of 0.01). **C, D)** Emetine and Imatinib combination results. **C)** Heatmap of the EOCDA, predicted minus observed combination viabilities, metric. Positive values (more red) suggest synergy whereas negative value suggest antagonism (more blue). **D)** Comparison of estimated and observed viabilities of Emetine and Imatinib in reference to the identity line (p-value = 0.392). **E, F)** GW 8510 and Withaferin A combination results. **E)** Heatmap of EOCDA. **F)** Comparison of estimated and observed viabilities (p-value = 0.27). **D, F)** Points are colored with the same scale as its corresponding EOCDA matrix. If the GoF p-value > 0.01 for the overall combination, then each point (i.e., dose) is classified as independent or non-independent drug action (See *Quantification of combinations not accounted for by the dCDA model* in Supplement).

To understand the extent to which the dCDA model captures the viability of cells in culture, we performed 26 sets of viability experiments under different doses and treatment times (see Methods). 84.6% of the combinations we studied were reasonable describable under the dCDA model (Fig. 2A X-labeled bars) of which a subset was also describable under the Bliss independence model (Fig. 2A double-dagger-labeled bars). We apply a Benjamini-Hochberg at an individual significance level of 0.05 to adjust for multiple hypotheses being tested. The subset of cases describable by Bliss independence corresponds to the dCDA model for *ρ* = 0. Equivalently, we can compute the difference between the estimated combination viability under the dCDA model and the observed combination viability, a metric we term the Excess over CDA (EOCDA). EOCDA is a measure of deviation from the dCDA model, and can be visualized as a heatmap where negative values (more blue) indicate an underestimation of the true viability and positive values (more red) indicate an overestimate of the true viability (Fig. 2C, E). If a combination is describable under dCDA, then the EOCDA metric also holds as an appropriate measure (Fig. 2B green and blue dots).

Classifying combinations as describable by the dCDA model or not is a global characterization and represents a measure across all of the combination viabilities at different doses taken together. We can be more granular and attempt to understand which specific doses in a dCDA compliant combination are likely synergistic or antagonistic. Let us first consider Fig. 2D and F where each (x,y) coordinate represents the (estimate, true) combination viability and are colored according to the EOCDA color scale. Possible antagonistic drug doses in this setup will be bluer points and possible synergistic drug doses will be redder points. Through a leave-one-out cross validation re-estimation framework (See *Quantification of combinations not accounted for by the dCDA model* in Supplement), we can assign each specific point (corresponding to specific doses of the two monotherapies) as locally following the dCDA model or not (p-value threshold of 0.01 for minimizing rate of falsely labelling points as not following the dCDA model). For instance, in Fig. 2D none of the points are locally likely synergistic or antagonistic but in Fig. 2F there are two such points (both likely synergistic). Of particular note, for the cases globally describable under dCDA presented in Fig. 2C, E and in Figs. S11 - S18, there are at most a few doses per combination that display locally likely synergistic or antagonistic behavior.

It is plausible, but not necessary, that the Spearman’s correlation estimate in the dCDA model describes to some extent the similarity between mechanisms of action of the individual drugs that constitute the combination. If that were the case, the larger the absolute value of the Spearman’s correlation estimate, the more similar the mechanisms of action are expected to be. Since our Spearman’s correlation estimates are on average not far from zero, we would expect that combinations describable under dCDA (GoF p-value > 0.002) are composed of monotherapies whose mechanisms of action are effectively unrelated. For example, Tamoxifen targets the estrogen receptor in breast cancers whereas GW 8510 is a cyclin-dependent kinase inhibitor investigated with respect to colorectal cancer [16, 17]. These two mechanisms of action are not related in any intuitive manner. This lack of a relationship is represented by a Spearman’s correlation estimate close to zero.

The majority of the tested combinations we studied were assayed at 24 hours after administering the drugs, but a select few were collected in a time-series experiment. There are two sets of combinations for which time-series viability data was collected. Experiments 21, 22, 23 correspond to Tamoxifen and Withaferin A at 12, 24, and 48 hours respectively (Fig. S18). The Spearman’s correlation estimate was stable over time but the EOCDA p-value decreased over time (0.68, 0.15, 3.3e-5). Thus, the dose CDA model sufficiently described the combination until the 48 hour time point. Experiments 5, 11, 26 correspond to Tamoxifen and Mefloquine at 12, 48, and 24 hours respectively (Fig. S17). The Spearman’s correlation estimates varied over time (0.12, -0.05, 0.02) but the EOCDA p-values were consistently above the threshold for rejection of the dCDA null model (p-value > 0.002). From both time-series experiments, we can observe that there may be undiscovered temporal dynamics at play with respect to the action of drug combinations in cell culture.

Independent of the aforementioned six experiments, we collected data for Tamoxifen and Mefloquine at 24 hours in Experiment 14 and data for Tamoxifen and Withaferin A at 24 hours in Experiment 12. As part of the 20 non-time series trials, we didn’t collect pure monotherapy viabilities but rather the viability at which the other drug in the combination was given in the lowest tested dosage. Thus, we can assess the robustness of the model to non-ideal input data by comparing Experiment 14 to Experiment 26 and Experiment 12 to Experiment 22. The Spearman’s estimate differs by 0.05 between Trial 14 and 26 and in both cases the dCDA model sufficiently describes the true data (Fig. 2A). The Spearman’s estimate differs by 0.01 between Trial 6 and 25 and once again, the dCDA model p-value is concordant between trials (Fig. 2A). These results favorably suggest that non-ideal input data does not considerably affect the performance of the model.

Excess over Bliss (EOB), one of the most popular performance metrics for assessing the synergistic or antagonistic behavior of a combination at particular dosages, is based on Bliss independence. In order for EOB to be a valid measure, the underlying combination must be describable under Bliss independence. Given that the dCDA model reduces to Bliss independence when *ρ* = 0, we can formally test whether the combination can be sufficiently explained by Bliss independence (See Methods). The EOB condition is valid for 50% (13 of the 26) of tested combinations (Fig. 2A, double-dagger-labelled bars). Intuitively, the closer the optimal Spearman’s correlation estimate is to zero, the more likely the EOB condition is to be valid. We can also investigate the joint distribution of p-value statistics for the EOB and EOCDA status of each tested combination (Fig. 2B). Most of the combinations tested can be described by dCDA (blue and green dots in Fig. 2B) whereas much fewer can be described by Bliss independence (blue and cyan dots in Fig. 2B). At the intersection, 38.5% of combinations are both describable under dCDA and satisfy Bliss independence (Fig. 2B, blue dots). It is important to note that, on the whole, the combinations describable under Bliss independence are a subset of those describable under dCDA. This highlights the strength of dCDA and EOCDA - we minimize our chances of identifying a combination as likely synergistic or antagonistic when it in fact explainable under independent but correlated joint action.

## Discussion

We introduced Correlated Drug Action (CDA) as a baseline model to explain combinations in the temporal and dose domains. The CDA model, based on simple principles, generalizes earlier models. Specifically, we discussed the tCDA model for the analysis of survival times in patient populations and the dCDA model for cell viability in cell cultures. We provided a statistical framework to assess the validity of the CDA which allows us to further classify drug combinations as likely synergistic and antagonistic, which is important to better understand combination therapies and to narrow down which combinations to investigate further.

As a model of survival times in patient populations treated with a drug combination, tCDA assumes [7, 8] that the monotherapies act independently in the population but that the times that each patient would survive under each monotherapies if delivered independently are correlated. Our tCDA formulation uses the same mathematical dependence as [8] to relate the effects of the single therapies to the survival under the combination and the potential correlation between survival times. This formula is proposed as an interpolation between two corner cases (correlation 0 and correlation 1) for which the mathematical expression can be found exactly. We show that that this model, derived as an interpolating approximation, actually represents the exact formula for a class of distributions of survival times, and that the correlation used in that formula should be the Spearman correlation. We studied the effect of different correlation structures between survival times and showed that the specific joint distribution of survival times does not substantially affect the results (Figs. S9). Therefore our results justify using the approximated analytical formula even when the underlying correlation structure between survival times is unknown (which is the case in general). Using an analytical formula for the tCDA rather than simulated joint distributions as done in [7] offers a computational advantage in terms of speed and efficiency in fitting the model to data, which allowed us to develop a statistical testing procedure to test the hypothesis that the actual survival in the combination follows the tCDA model. We show that the tCDA model accurately describes a major subset of the clinically relevant combination cancer therapies remarkably well (Fig. 1A, S1).

The public data collected in this work to validate the tCDA model uses data from 18 clinical trials, which include nine that were described in [7]. In order to collect clinical trial data that was in the format necessary for the tCDA model, we often had to aggregate data from up to three different clinical trials (Supp. Table 3). In these scenarios, we attempted to match cohort characteristics including sex, age, ethnicity, and previous treatment history as well as possible.

The tCDA model predicts the distribution of survival times for cohorts treated with combination therapy. To generalize these ideas to predict cell viability in cell cultures treated with drug combinations at different doses of the constituent drugs, we introduced the dCDA model. The dCDA model generalizes popular reference models proposed in the literature. A key and novel feature in the theory leading to the dCDA model is the assumption that for each cell in culture treated with a drug for a certain time T, there exists a lethal dose (dependent on T and varying from cell-to-cell) such that, if the drug is applied at a dose lower (resp. higher) than the lethal dose, the cell will be found alive (resp. dead) after the given treatment time T. If two drugs are applied for a time T in combination and at their respective doses, the dCDA model postulates that the only cells that survive are those for which both lethal doses are higher than the doses at which the drugs in the combination were applied. Note that the lethal dose for each cell in a culture in the dCDA context plays a similar role to the survival time for each patient in a population in the tCDA context. Like the tCDA model, the dCDA model can be formulated as an exact closed-form expression if we assume a particular joint distribution of lethal doses in the cell population, parameterized by Spearman’s correlation *ρ* between lethal doses. As the true joint distribution of lethal doses is actually unknown, and the results are insensitive to the specific joint distribution, we used the closed formula to analyze experimental data. The dCDA model reduces to the the Bliss independence model for *ρ* = 0 and, by leveraging the Dose Equivalence Principle, to sham-compliant HSA for *ρ* = 1 (See *Formulation of dose-space correlated drug action model* in Supplement). This defines a spectrum of models wherein varying the Spearman’s correlation from zero to one controls whether the combination is more Bliss-like or more sham compliant HSA-like. The actual value of *ρ* is determined by the data, and estimated by our framework (Fig. 2 and Supp. Table 2). Thus our paradigm combines the advantages of several ways of thinking about drug interaction in the dose space.

To discover likely synergistic/antogonistic interactions between drugs, we introduced a new metric which we called Excess over Correlated Drug Action, or EOCDA, which is simply the difference between the viability estimated using the dCDA model and the observed viability; positive EOCDA values correspond to likely synergistic combinations and negative EOCDA values correspond to likely antagonistic combinations (Fig. 2C,E). We compared the EOCDA metric to the popular Excess over Bliss (EOB) metric. For both the EOCDA and EOB metrics we defined a statistical significance test to determine if the null hypothesis (EOCDA=0 or EOB=0) should be rejected. Therefore, for each of the tested combinations, we have a p-value for the null hypothesis EOB = 0 and EOCDA = 0. Fig. 2B illustrates the degree of relatedness between the two metrics. We can notice that there are more cases for which we cannot reject the null hypothesis of independent drug action (at a significance level of 0.01) under the dCDA when the real viability is compared to the viability estimated using Bliss independence (more blue and green dots than blue and yellow dots in Fig. 2B). In other words, EOCDA reduces the chance of calling a combination likely synergistic or antagonistic compared to EOB - an important criteria when considering the experimental resources and effort that it takes to validate effective drug combinations. One limitation of EOCDA, however, is that we need the entire dose matrix with corresponding viabilities to estimate the parameter *ρ* whereas EOB can be computed at individual dosages. Interestingly, for combinations describable under dCDA, we observe that there are only a few doses that are “locally” likely synergistic or antagonistic combinations in a globally independent-drug combination. This plausibly identifies dosages of particular interest for further investigation.

As discussed earlier, the fitted parameter *ρ* in the dCDA corresponds to the Spearman’s rank correlation between the lethal doses of two different therapies for each cell across the cell culture. The Spearman’s correlation provides us information regarding the association between the two monotherapies including the possibility of the two drugs have similar mechanisms of action. The reason we are able to discuss possible biological explanation from a much more general definition in a cell culture case is that unlike patients, cell cultures are relatively homogeneous and isogenic tumor cells free of the external milieu and confounders that exists in a human population. In our data, all of the correlation estimates are weak implying that, from the perspective of dCDA, the drugs probably act using alternative pathways (Fig. 2A). For example, GW 8510 is a cyclin-dependent kinase inhibitor whereas Withaferin A inhibits Vimentin, an intermediate filament. These two mechanisms are not linked by any known direct relationship, and this supports the low Spearman’s correlation estimate given by the dCDA model (Fig. 2E, F). More work is needed to validate the notion that similarity in the mechanism of action of two drugs can be reflected in the value of the parameter *ρ*.

There are many possible avenues for extending this work. One such extension could be the creation of a network of therapies for a given disease. Each node is a particular monotherapy and the edge width between two nodes corresponds to Spearman’s correlation estimate. This could be done for cell cultures or patient populations. From this setup, we can attempt to learn on the graph and infer the edge strength between other pairs of nodes. This could be informative in suggesting new candidates for cell line experiments or clinical trials. Additionally, we can investigate the relationship between the dose-space and temporal CDA models by testing the same cancer combination therapy in mice PDX models and patient cancer cells cultured in the lab. Finally, a more general theory that combines both dose models and temporal models into one single model seems plausible and potentially useful.

## Methods

### Collection of clinical trial data

We collected data from clinical trials using the clinicaltrials.gov query tool and scraped Progression Free Survival (PFS) data from their associated results and papers. We converted graphs into tables of values through a robust online digitizer (link). Though it would be best to have the original data, these files were not available to us. For a given combination, we aimed to find a single trial that tested the individual therapies and the combination. However, we also had to pull data across different trials to construct the necessary survival data. In these cases, we tried to maximize overlap in patient characteristics, dosages, and time schedules. We were able to match dosages and time schedules in the overwhelming majority of cases. We assembled data for 18 distinct combinations spanning 10 different cancer types. A detailed guide to the raw data can be found in Supp. Table 3. The tables of the raw PFS curves from the digitized plots that we used for our analysis can be found in Supp. File 1.

### tCDA model

Code for the Temporal CDA Model, tCDA, can be found [here](https://github.com/aditharun/correlated-drug-action/blob/main/scripts/clinicaltrial.R). Results for tCDA on the clinical trials can be found in Supp. Table 1.

### Drug Combination Experiments in the MCF7 cell line

#### Choice of Drug Combinations, Dose matrix and Treatment Times

We applied the dCDA model to fit the viability of MCF7 cells in response to 26 drug combinations. These 26 drug combinations comprised 20 unique combinations, and 6 repeat combinations as discussed below. The 20 combinations, which included 13 unique drugs (Atovaquone, Emetine, GW 8510, Imatinib, Mefloquine, MG 132, Mitoxanthrone, Nicardipine, Sanguinarine, Tamoxifen, Terfenadine, Tyrphostin AG 825 and Withaferin A), were selected from a a pre-existing drug combination screen collected at Columbia University (Andrea Califano lab) in the context of the Library of Integrated Network-Based Cellular Signatures (LINCS) Program. This screen tested all 990 combinations of 99 drugs against 10 drugs, each combination assessed in a dose response matrix of 4×4 doses. Of these combinations we chose a subset of combinations that exhibited antagonistic, independent and synergistic behaviors, resulting in the 20 combinations reported here. The 20 drug combinations were administered at doses shown in the 6×6 dose matrices shown in Figs. 2, and Figs. S10 - S15, and the viability was measured at 24 hours. These 20 experiments, which we refer to as the 24h viability experiments, are reported for the first time in this paper and provided as Supp. File 2. Amongst the 20 drug pairs, two specific combinations (Tamoxifen/Withaferin A and Tamoxifen/Mefloquine) were measured at more detailed dose matrices and treatment times as reported in an earlier publication [18]. For these two pairs, viability was measured at the doses corresponding to the 10×10 dose matrix shown in Figs. S16 and S17, and at treatment times of 12h, 24h and 48h. We also measured the viability of the individual monotherapies (Tamoxifen and Mefloquine) at the 10 doses reported in the figures, at 12h, 24h, and 48h. These 6 experiments, which we will refer to as the time course viability experiments, had been previously made publicly available in [18]; we also provide these data for convenience in Supp. File 2. All drug combination viability experiments (the 24h and time course experiments) were done in triplicate and the results for each combination were averaged. All monotherapy experiments (for the time course data) were done in duplicate and the results averaged. The dCDA model requires that we have viability for the monotherapies. We had that data for the time course data, but we lacked the equivalent data for the 24h viability experiments. Therefore to apply the dCDA model, we approximated the cell viability under individual drugs by considering the case when the other drug in the combination was measured at its lowest concentration.

#### Cell Culture

MCF7 (ATCC HTB-22) cells were obtained from ATCC. Cells were cultured according to manufacturer’s recommendations in ATCC-formulated Eagle’s Minimum Essential Medium (Catalog No. 30–2003) with 10% heat-inactivated fetal bovine serum, and 0.01 mg/ml human recombinant insulin. Growth media was changed every 3–4 days. After reaching confluence, cells were split at a ratio 1:6. Cultures were tested for mycoplasma periodically using MycoAlert (Lonza, Cat No. LT07-701) per manufacturer’s instructions. To split, media was removed, cells were washed with PBS, and trypsin-EDTA mix was added for 5 min. After detachment, cells were washed with growth media, collected into 50 ml vial, spin down at 1000 RPM, suspended in fresh media and plated into 75 cm flasks. The cells were plated at 10,000 cells per well in a clear bottom black 96-well plate (Greiner Cat. No. 655090) and a white 96-well plate (Greiner Cat. No. 655083) then they were placed in an incubator. After 24 hr, the plates were removed from the incubator and treated with drugs using the HP D300 Digital Dispenser.

#### Cell viability

Cells were then treated with the appropriate drug combinations at specific dose matrices and treatment times. After the targeted drug treatment time, 100 μL of Cell-Titer-Glo (Promega Corp.) was added to the wells in the white 96-well plate and shaken at 500 rpm for 5 min. The plate was then read by the Perkin Elmer Envision 2104 using an enhanced luminescence protocol to count the number of raw luminescent units per well. For each viability experiment, the results were normalized by the luminescence measured in untreated cell cultures. The ratio of luminescence in treated vs untreated cells is what we report as cell viability.

### dCDA model

Code for the dose-space CDA model, dCDA, can be found here here. Results for dCDA on the experimental combinations can be found in Supp. Table 2.

## Acknowledgements

We thank the Columbia University High Throughput Screening Facility and the Sulzberger Columbia Genome Center, where the viability experiments were conducted by Charles Karan and Ronald B. Realubit. We also thank Andrea Califano for providing the LINCS screen of 990 drug combinations from where we chose the drug combinations for cell viability used in this paper.

## Authors’ contributions

GS conceived of the project. All co-authors developed the methodology for the project. ASA and SCK developed computational approaches and performed bioinformatics analysis with guidance from GS and MEA. ASA drafted the manuscript and all authors participated in its editing and revision.

## Availability of Data and Materials

All data and code can be found at github.com/aditharun/correlated-drug-action.

