## Supplemental Text for "Correlated Drug Action as a Baseline Model for Combination Therapy in Patient Cohorts and Cell Cultures"

### Temporal Correlated Drug Action

Temporal Correlated Drug Action (tCDA) is an independent joint action model for the action of drug combination on patients when studying their survival times. Let us assume that the effect of a therapy on a patient population can be evaluated using the distribution of survival times such as Progression Free Survival (PFS) or Overall Survival (OS). tCDA postulates that the effect of two therapies in combination is the same as the effect of the single best therapy if they had been given independently. In other words, if we knew a priori which of two therapies yields better results for a given patient (something that precision medicine can not predict today), tCDA postulates that the effect of administering the best monotherapy for that patient would be the same as the effect of administering the combination.

The survival times may vary from patient to patient due to different individual conditions such as tumor composition, immune system status and co-morbidities among others reasons. tCDA assumes no direct interaction between the two therapies administered to a patient. However, the hypothetical survival times on the same patient due to each of two drugs could be correlated due to the fact that the individual patient could act as a confounding factor through either the nature of the tumor, the age, the health status, etc. For example, a patient with an aggressive tumor would have a shorter survival if treated with either monotherapy, while a patient with a less aggressive tumor would survive longer with either monotherapy. In this case, the survival times under each monotherapy will be correlated.

The tCDA framework provides a baseline hypothesis for independent action of the drugs that can be used as a null hypothesis in a formal hypothesis test to classify clinical combinations as consistent with tCDA or inconsistent with tCDA. Non-tCDA combinations that are of particular interest are antagonistic or synergistic combinations.

#### Notation

For a patient in a cohort, we will denote by $t_A$, $t_B$ and $t_{AB}$ the times that the patient would survive if treated with drug A, drug B or the combination of A and B respectively. Under the tCDA model, the survival time $t_{AB} = \max(t_A, t_B)$ for a given patient. Note that in practice we can only measure one of these survival times as a patient will only be treated with either treatment, and their survival will only be registered under that treatment. However, we can computationally simulate the survival times under the other treatment for the purposes of developing the tCDA model.

The survival probability that a generic patient survives a time $t$ under treatments A, B and combination AB will be denoted by $S_A(t)$, $S_B(t)$ and $S_{AB}(t)$, and can be formally expressed as

$$S_A(t) = \text{Prob}(t_A > t) \quad (1)$$

$$S_B(t) = \text{Prob}(t_B > t) \quad (2)$$

$$S_{AB}(t) = \text{Prob}(\max(t_A, t_B) > t) \quad (3)$$

$$\quad (4)$$

In practice the probabilities are computed as fractions in a cohort. For example, the $S_A(t)$ is the fraction of patients in a cohort whose survival time was longer than $t$.

The hypothetical survival times $t_A$ and $t_B$ that a patient would survive under drugs A and B will in general be correlated through confounding factors. We will consider two types of correlations between these times. The Pearson correlation will be denoted by $\rho_P$ and the Spearman's correlation, the correlation between the ranks, will be denoted by $\rho_s$.

#### Formulation of the Temporal Correlated Drug Action Model

In this section we want to derive expressions for the survival probability that a patient has a survival time equal to or longer than $t$ under treatment with both drugs A and B in terms of the survival probabilities under the monotherapies. Under the tCDA model, if a patient survives more than time $t$ under the AB combination, we have either $t_A > t$ or $t_B > t$. Therefore, from (3)

$$S_{AB}(t) = \text{Prob}(t_A > t \text{ OR } t_B > t) \tag{5}$$

$$= 1 - \text{Prob}(t_A \le t \text{ AND } t_B \le t), \tag{6}$$

where we used that the complement of a union ("OR" logical operation) of two sets is the intersection ("AND" logical operation) of the complement of the two sets. To proceed we consider separately the correlated and the uncorrelated cases.

##### Uncorrelated Case

We will consider the case where the two drugs A and B are uncorrelated because they are statistically independent from one another. (In the general case, the drugs could be uncorrelated but statistically dependent). In this case, Prob$(t_A \le t$ AND $t_B \le t) =$ Prob$(t_A \le t)$Prob$(t_B \le t)$, which leads to

$$S_{AB}(t, \rho_s = 0) = S_A(t) + S_B(t) - S_A(t)S_B(t), \tag{7}$$

where we used that Prob$(t_A \le t) = 1 - S_A(t)$. Note that in this case, the survival under the two drugs is greater than the survival under either of the single therapies. Note the similarity between this formula and the equivalent formula for Bliss independence when computing the inhibition of a cell culture measured at a particular time after application of a drug combination. In that case the inhibition of the cell culture under the Bliss independence model is estimated as $I_{AB} = I_A + I_B - I_A I_B$.

##### Positively correlated case

Let us consider the extreme case in which for any given patient the survival time $t_A$ completely determines the survival time $t_B$, in such a way that the Spearman's correlation between survival times under A and B treatments of all patients is 1. That means that the order of survival times of patients under treatment A is the same as the order under treatment B. Take now a patient with survival times $t_A$ and $t_B$. The fraction $S_A(t_A)$ of patients that survived more than the patient under A, should be the same as the fraction $S_B(t_B)$ of patients that survived more than the patient under B. Therefore when the Spearman's correlation between survival times is $\rho_s = 1$, we have that

$$t_B = S_B^{-1}(S_A(t_A)). \tag{8}$$

In this extreme case we can compute the survival time under the combination AB. First we note that we can express Prob($t_A \leq t$ AND $t_B \leq t$) in terms of conditional probabilities as

$$\begin{aligned}\text{Prob}(t_A \leq t \text{ AND } t_B \leq t) &= \text{Prob}(t_A \leq t | t_B \leq t)\text{Prob}(t_B \leq t) &(9)\\ &= \text{Prob}(t_B \leq t | t_A \leq t)\text{Prob}(t_A \leq t). &(10)\end{aligned}$$

Suppose now that $S_B(t) > S_A(t)$. This means that at time $t$, drug B is better than A, in that there are more patients that survived with drug B more than time $t$ than with drug A. Because the order of survival of the patients is the same under both drugs then all the patients that did not survive under B, couldn't have survived under A. Therefore if $B$ is better than $A$ Prob($t_A \leq t | t_B \leq t) = 1$. Symmetrically, if $A$ is better than $B$ Prob($t_B \leq t | t_A \leq t) = 1$.

On the other hand, when $t_A$ and $t_B$ and uncorrelated and independent, we have that Prob($t_A \leq t | t_B \leq t$) = Prob($t_A \leq t$) and symmetrically, Prob($t_B \leq t | t_A \leq t$) = Prob($t_B \leq t$). In the intermediate case between in which we have some correlation $\rho_s$, we will interpolate the conditional probabilities in such a way that the completely correlated and uncorrelated case case match the extreme cases as follows

$$\text{Prob}(t_A \leq t | t_B \leq t) = \alpha(\rho_s) + (1-\alpha(\rho_s))\text{Prob}(t_A \leq t) \text{ if } S_B(t) > S_A(t) \quad (11)$$

$$\text{Prob}(t_B \leq t | t_A \leq t) = \alpha(\rho_s) + (1-\alpha(\rho_s))\text{Prob}(t_B \leq t) \text{ if } S_B(t) \leq S_A(t) \quad (12)$$

where $\alpha(\rho_s = 0) = 0$, $\alpha(\rho_s = 1) = 1$ and the specific functional form in the range $0 \leq \rho_s \leq 1$ will depend on the joint distribution of survival times.

Replacing these results in (10) and using (3) we find the survival under the combination AB in the case of $0 \leq \rho_s \leq 1$ is

$$S_{AB}(t, 0 \leq \rho_s \leq 1) = \begin{cases} S_A(t) + S_B(t)(1 - S_A(t))(1-\alpha(\rho_s)) & \text{if } S_A(t) > S_B(t) \\ S_B(t) + S_A(t)(1 - S_B(t))(1-\alpha(\rho_s)) & \text{if } S_A(t) \leq S_B(t) \end{cases} \quad (13)$$

**Negatively correlated case**

Let us consider the other extreme case in which for any given patient the survival t ime $t_A$ completely determines the survival time $t_B$, in such a way that the Spearman correlation, $\rho_s$, between survival times under A and B treatments of all patients is $-1$. That means that the order of survival times of patients under treatment A is the opposite as the order under treatment B. Take now a patient with survival times $t_A$ and $t_B$. Therefore, $S_A(t_A)$ represents the fraction of patients that survived more than the patient in question under treatment A; this should be the same as $1 - S_B(t_B)$, the fraction of patients that died earlier than the patient in question under B. Thus, when the Spearman correlation between survival times is $\rho_s = -1$, we have that

$$t_B = S_B^{-1}(1 - S_A(t_A)). \quad (14)$$

In this extreme case we can compute the survival time under the combination AB. First we note that there is a time $t^*$ such that all the patients that lived more than $t^*$ under A lived less than $t^*$ under B. This condition is given by $S_A(t^*) = 1 - S_B(t^*)$. Figure S1 shows that for $t > t^*$, the fraction of patients that survived more than $t$ under A or B is the sum of those who survived more than $t$ under A and those who survived more than $t$ under B, given that the anticorrelation makes these groups non-overlapping.

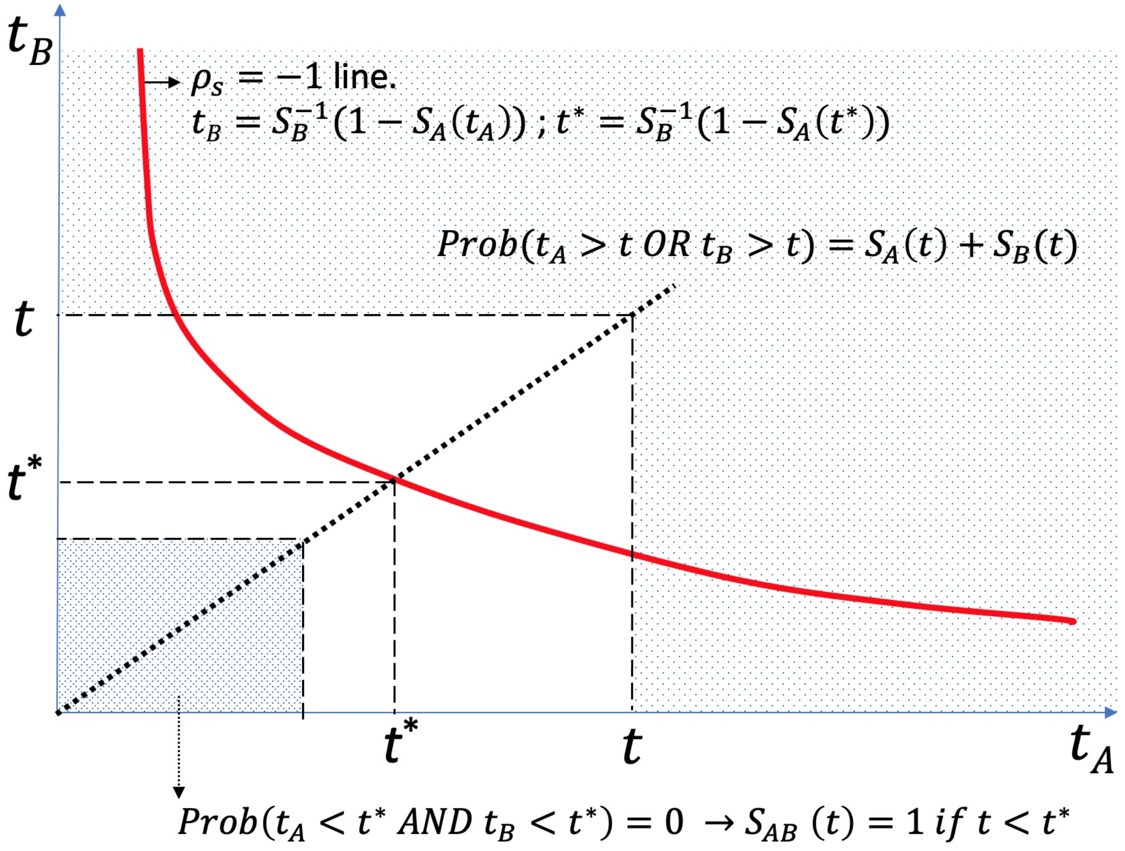


**Figure S1: Case $\rho_s = -1$.**

Figure S1 also shows that for $t < t*$, all patients survived more than $t$ thanks to either monotherapy or both monotherapies. Therefore for $\rho_s = -1$, we have that

$$S_{AB}(t, \rho_s = -1) = \begin{cases} 1 & \text{if } t \leq t^* \\ S_A(t) + S_B(t) & \text{if } t > t^* \end{cases} \tag{15}$$

where $t^*$ is defined as the time that makes $S_A(t^*) + S_B(t^*) = 1$.

In the range of negative correlations, we can interpolate between the cases of $\rho_s = 0$ and $\rho_s = -1$, which yields

$$S_{AB}(t, -1 \leq \rho_s \leq 0) = \begin{cases} [S_A(t) + S_B(t) - S_A(t)S_B(t)][1 - \alpha(\rho_s)] + \alpha(\rho_s) & \text{if } t \leq t^* \\ S_A(t) + S_B(t) - S_A(t)S_B(t)[1 - \alpha(\rho_s)] & \text{if } t > t^* \end{cases} \tag{16}$$

where $\alpha(\rho_s = 0) = 0$, $\alpha(\rho_s = -1) = -1$, and the specific dependence of $\alpha(\rho_s)$ on $\rho_s$ depends in the joint distributions of times $t_A$ and $t_B$.

##### Integrated expression for the survival probability under the tCDA model

The formulas of the previous subsection can be integrated in one simple formula. Call $S_{\text{max}}(t) = \max(S_A(t), S_B(t))$, and $S_{\text{min}}(t) = \min(S_A(t), S_B(t))$. The expression for the survival probability under the tCDA model can be written as:

$$S_{AB}(t, \rho_s) = \tag{17}$$
$$= \begin{cases} S_{\text{max}}(t) + S_{\text{min}}(t)(1 - S_{\text{max}}(t))(1 - \alpha(\rho_s)) & \text{if } 0 \leq \rho_s \leq 1 \\ [S_A(t) + S_B(t) - S_A(t)S_B(t)][1 - \alpha(\rho_s)] + \min(1, S_A(t) + S_B(t))\alpha(\rho_s) & \text{if } -1 \leq \rho_s \leq 0 \end{cases}$$

The model is not complete until we specify the function $\alpha(\rho_s)$, which is unknown except in the border cases $\rho_s$ =-1, 0, 1 for which $\alpha(\rho_s)$= is -1, 0, 1 respectively. As an approximation, we can assume that $\alpha(\rho_s) = \rho_s$. In the next section we will show that this is indeed the correct function for a particular joint distribution for $t_A$ and $t_B$.

##### There is a joint survival times distribution for which $\alpha(\rho_s) = |\rho_s|$

There are many possible joint distributions of $t_A$ and $t_B$ that yield the same Spearman correlation, and for each such distribution there may be a different function $\alpha(\rho_s)$. Therefore from the perspective of defining uniquely the tCDA model, it is necessary to specify the joint distribution of survival times. In this section we will choose such distribution and show that for it, $\alpha(\rho_s) = \rho_s$, which we will call the tCDA joint survival times probability density, or tCDA distribution for short.

This distribution can be thought as a mixture distribution where a fraction $0 \leq \alpha \leq 1$ of ($t_A$, $t_B$) pairs are drawn from the line $t_B = S_B^{-1}(S_A(t_A))$ for $0 \leq \rho_s \leq 1$ or the line $t_B = S_B^{-1}(1 - S_A(t_A))$ for $0 \leq \rho_s \leq 1$, and the other $1 - \alpha$ fraction are independently drawn from the distributions $P_A(t_A)$ and $P_B(t_B)$. Formally this can be written as

$$P(t_A, t_B) = (1 - \alpha)P_A(t_A)P_B(t_B) + \alpha \begin{cases} \delta(S_A(t_A) - S_B(t_B))P_A(t_A)P_B(t_B) & \text{if } 0 \leq \rho_s \leq 1 \\ \delta(S_A(t_A) + S_B(t_B) - 1)P_A(t_A)P_B(t_B) & \text{if -1} \leq \rho_s \leq 0 \end{cases} \tag{18}$$

Where $\delta(x)$ is the Dirac's Delta function. Note that, if $X$ denotes either A or B, $S_X(t_X) = \int_{t_X}^{\infty} P_X(t)dt$ and $D_X$ is the cumulative distribution of $P_X(t)$, then $S_X(t) = 1 - D_X(t)$. It is not difficult to show that the marginal distributions of $P(t_A, t_B)$ coincide with $P_A(t_A)$ and $P_B(t_B)$. We will now use (6), for which we need to integrate (18) in the rectangle $0 \leq t_A \leq 1$ and $0 \leq t_B \leq 1$. The term pertaining to the fraction $1 - \alpha$ of independently chosen points $t_A$ and $t_B$ can be easily integrated to yield

$$\int_{t_A=0}^{t} \int_{t_B=0}^{t} P_A(t_A)P_B(t_B)dt_A dt_B = (1 - S_A(t))(1 - S_B(t)). \tag{19}$$

To handle the second term we have to consider the positive and negative correlation cases independently.

*1) Case* $0 \leq \rho_s \leq 1$

We change variables to $z_A = S_A(t_A)$ and $z_B = S_B(t_B)$.

$$\int_{t_A=0}^{t}\int_{t_B=0}^{t} \delta(S_A(t_A) - S_B(t_B))\ P_A(t_A)P_B(t_B)dt_A dt_B =$$
$$= \int_{S_A(t)}^{1}\int_{S_B(t)}^{1} \delta(z_A - z_B)dz_A dz_B = \int_{S_{max}(t)}^{1} dz_{max} = 1 - S_{max}(t), \tag{20}$$

where we first integrated between the variable $z_{min}$ such that $S_{min}(t) = \min(S_A(t), S_B(t))$ to ensure that the first integral yielded 1. Putting together (19) and (20) and using (6) we get

$$\begin{aligned} S_{AB}(t) &= 1 - \alpha(1 - S_{max}(t)) + (1-\alpha)(1 - S_{max}(t))(1 - S_{min}(t)) \\ &= S_{max}(t) + S_{max}(t)(1 - S_{min}(t))(1-\alpha). \end{aligned} \tag{21}$$

which coincides with the positive $\rho_s$ case of (17).

*2) Case* $-1 \leq \rho_s \leq 0$
Starting from the negative $\rho_s$ branch of (18) we change variables to $z_A = S_A(t_A)$ and $z_B = 1 - S_B(t_B)$.

$$\begin{aligned} \int_{t_A=0}^{t}\int_{t_B=0}^{t} &\delta(S_A(t_A) + S_B(t_B) - 1)\ P_A(t_A)P_B(t_B)dt_A dt_B = \\ &= \int_{S_A(t)}^{1}\int_{0}^{1-S_B(t)} \delta(z_A - z_B)dz_A dz_B \\ &= \int_{S_A(t)}^{1}\int_{0}^{S_A(t)} \delta(z_A - z_B)dz_A dz_B + \int_{S_A(t)}^{1}\int_{S_A(t)}^{1-S_B(t)} \delta(z_A - z_B)dz_A \\ &= \max(0, 1 - S_A(t) - S_B(t)) \end{aligned} \tag{22}$$

where we put together (19) and (22) and using (6) we get

$$\begin{aligned} S_{AB}(t) &= 1 - \alpha \max(0, 1 - S_A(t) - S_B(t)) + (1-\alpha)(1 - S_A(t) - S_B(t) + S_A(t)S_B(t)) \\ &= \alpha \min(1, S_A(t) + S_B(t)) + (1-\alpha)(S_A(t) + S_B(t) - S_A(t)S_B(t)) \end{aligned} \tag{23}$$

which coincides with the negative $\rho_s$ case of (17).

To see the relation between the Spearman's correlation $\rho_s$ and the parameter $\alpha$, we consider a cohort of $N$ patients, each one with a pair of survival times $t_A$ and $t_B$. For each sample we will have the ranking of the survival times which we will call $i_A$ and $i_B$. Clearly $1 \leq i_A, i_B \leq N$.The probability over all possible cohorts of having a patient with a pair of survival times ($t_A$, $t_B$) will be denoted by $\mathcal{P}(i_A, i_B)$. As before, we will consider a mixture distribution with a fraction $1-\alpha$ of the patients with random ranks $i_A$ and $i_B$ (corresponding to a distribution $\mathcal{P}_r(i_A, i_B) = \frac{1}{N^2}$, and the other fraction of $\alpha$ patients being either perfectly correlated (with distribution $\mathcal{P}_{\rho_s=1}(i_A, i_B) = \frac{1}{N}\delta_{i_A,i_B}$) or perfectly anti-correlated (with distribution $\mathcal{P}_{\rho_s=1}(i_A, i_B) = \frac{1}{N}\delta_{N+1-i_A,i_B}$),

$$\mathcal{P}(i_A, i_B; \rho_s) = (1-\alpha)\frac{1}{N^2} + \alpha \begin{cases} \frac{1}{N}\delta_{i_A,i_B} & \text{if } 0 \leq \rho_s \leq 1 \\ \frac{1}{N}\delta_{N+1-i_A,i_B} & \text{if } -1 \leq \rho_s \leq 0 \end{cases} \tag{24}$$

The Spearman correlation is the rank correlation of the survival times, and as such, it can be computed as

$$\rho_s = \frac{\langle i_A i_B \rangle - \langle i_A \rangle \langle i_B \rangle}{\sigma_{i_A} \sigma_{i_B}}. \tag{25}$$

where $\delta_{i,j}$ is Kronecker's delta, $\sigma^2_{i_A} = \langle i_A^2 \rangle - \langle i_A \rangle^2$ and $\sigma^2_{i_B} = \langle i_B^2 \rangle - \langle i_B \rangle^2$. To proceed, we compute $\langle i_A i_B \rangle$, $\langle i_A \rangle$ and $\langle i_B \rangle$ using the expression of $\mathcal{P}(i_A, i_B; \rho_s)$ for positive and negative $\rho_s$. Note that the marginal distributions of $i_A$ and $i_B$ are the same regardless of the sign of $\rho_s$, that is $\mathcal{P}(i_B) = \frac{1}{N}$, and therefore

$$\langle i \rangle := \langle i_A \rangle = \langle i_B \rangle = \frac{N+1}{2} \tag{26}$$

$$\langle i^2 \rangle := \langle i_A^2 \rangle = \langle i_B^2 \rangle = \frac{(N+1)(2N+1)}{6} \tag{27}$$

$$\sigma_i^2 := \sigma^2_{i_A} = \sigma^2_{i_B} = \frac{(N+1)(N-1)}{12} \tag{28}$$

*i) Case positive $\rho_s$:*

In this case

$$\langle i_A i_B \rangle = (1-\alpha)\frac{1}{N^2} \sum_{i_A=1}^{N} i_A \sum_{i_B=1}^{N} i_B + \alpha \frac{1}{N} \sum_{i_A=1}^{N} i_A \sum_{i_B=1}^{N} i_B \delta_{i_A, i_B} \tag{29}$$

$$= (1-\alpha)(\frac{1}{N} \sum_{i=1}^{N} i)^2 + \alpha \frac{1}{N} \sum_{i=1}^{N} i^2 \tag{30}$$

$$= (1-\alpha)\langle i \rangle^2 + \alpha \langle i^2 \rangle \tag{31}$$

Using (25) we find that

$$\rho_s = \frac{(1-\alpha)\langle i \rangle^2 + \alpha \langle i^2 \rangle - \langle i \rangle^2}{\sigma_i^2} = \alpha \frac{\langle i^2 \rangle - \langle i \rangle^2}{\sigma_i^2}$$

$$= \alpha. \tag{32}$$

*ii) Case negative $\rho_s$:*

In this case

$$\langle i_A i_B \rangle = (1-\alpha)\frac{1}{N^2} \sum_{i_A}^{N} i_A \sum_{i_B=1}^{N} i_B + \alpha \frac{1}{N} \sum_{i_A=1}^{N} i_A \sum_{i_B=1}^{N} i_B \delta_{N+1-i_A, i_B} \tag{33}$$

$$= (1-\alpha)(\frac{1}{N} \sum_{i=1}^{N} i)^2 + \alpha \frac{1}{N} \sum_{i=1}^{N} (N+1-i) i \tag{34}$$

$$= (1-\alpha)\langle i \rangle^2 + \alpha (\frac{(N+1)^2}{2} - \langle i^2 \rangle)$$

$$= (1+\alpha)\langle i \rangle^2 - \alpha \langle i^2 \rangle \tag{35}$$

Using (25) we find that

$$\rho_s = \frac{(1+\alpha)\langle i\rangle^2 - \alpha\langle i^2\rangle - \langle i\rangle^2}{\sigma_i^2} = \alpha\frac{\langle i\rangle^2 - \langle i^2\rangle}{\sigma_i^2}$$
$$= -\alpha. \tag{36}$$

It follows that for this choice of distribution, the tCDA model is exact and corresponds to $\alpha(\rho_s) = |\rho_s|$.

#### Pairing survival times under two drugs to attain a given Spearman correlation

Given two survival curves corresponding to two monotherapies A and B that are also tested in combination, we would like to have a method to pair samples of survival times in such a way that we have a given Spearman's correlation. This can be done by drawing $N$ independent samples $t_A$ consistent with drug A's survival curve (note: 1 - survival curve = cumulative distribution). Likewise, we can draw another set of $N$ independent samples $t_B$ consistent with drug B's survival curve and pair these survival times until they have a desired Spearman correlation.

There are many possible ways to reorganize two vectors such that they have a specified correlation while keeping the corresponding marginal distributions. Below we discuss two different ways to accomplish this, but there are many other ways to do it that we have not explored. We believe that each method for randomizing with correlation is equally likely to be the true method by which nature operates, but we are unaware of any way to discern which method may be more likely to be true. Thus, we proceed with the two methods introduced here. One is the numerical implementation of the theoretical joint distribution discussed in the previous section, and as such, it has the advantage that we can use analytical expressions for their implementation. The second method is used as a comparative alternative, to determine the dependence of the results on the specific way of pairing the survival times. We will see that the final conclusions are robust to the specific pairing method used. Both methods start by fixing one vector of times, say $t_A$, and shuffle the other vector appropriately until the specified correlation is reached to within a tolerance.

*The window-swap method*
Let us assume that our target correlation $\rho_s$ is positive. The method starts with two vectors of times $t_{A,i}$ and $t_{B,i}$ ordered from the minimum values $t_{A,1}$ and $t_{B,1}$, to their maximum values $t_{A,N}$ and $t_{B,N}$, in such a way that their Spearman correlation is 1. To attain our target Spearman correlation we "disorganize" the relative order of the $t_A$ and $t_B$ vectors in a controlled way. To do that we go through each index $i$ and use a symmetric window of size $w$ around the current index comprising indices ranging from $i-w$ to $i+w$ (if $i-w$ is less than 1, we clamp the minimum at 1; if $i+w$ is larger than $N$, we clamp the maximum at $N$). We then choose a random index $j$ in the interval $[i=w, i+w]$ and swap the values of $t_B$ at positions $j$ and $i$. After visiting all indices we compute the Spearman correlation. If its value is larger than the target one, we restart the process with a larger $w$ (e.g., doubling the previous one, or increasing the previous one to $w+1$). If the last iteration created a Spearman correlation that is smaller than the target, we go back to the previous iteration and redo the last iteration with a smaller window (e.g. halving the previous one or increasing the previous one to $w-1$). This process converges when we arrive at a Spearman correlation that differs from the target correlation within a pre-specified tolerance, e.g., 1%. The previous process need only work for positive target correlation values. If our target correlation $\rho_s$ is negative, we find the shuffled vector using the absolute value of $\rho_s$ and once

it converges, we simply flip the shuffled $t_B$ vector to get $t'_B$, the appropriate vector (i.e. reverse order such that $t'_{B,i} = t_{B,N+1-i}$).

*The coin method*
The coin method is parametrized by the weight of the coin - the probability $\alpha$ of landing heads. Similar to the window-swap method, this method starts with two vectors of times $t_{A,i}$ and $t_{B,i}$ ordered from the minimum to the maximum values. For each index $i$ in the to-be-shuffled vector, we flip the weighted coin with weight $\alpha$. If heads, we randomly swap the current value $t_{B,i}$ with the value $t_{B,j}$ at an index $j$ randomly chosen in the interval $[1, N]$ but excluding the current index $i$. This coin method of simulation is an implementation of the theory presented in the previous section. Therefore the target Spearman correlation, $\rho_s$, can directly be used as the weight of the coin according to $\alpha = |\rho_s|$. Just as for the window swap method, we flip the resulted shuffled vector if the desired Spearman's correlation is negative. Fig. S7 exemplifies the results of the previous section in the context of the coin method. Fig. S7A shows the results of the coin method to create a joint distribution of survival times for which a target fraction $|\rho|$ of points falls on the orange line (the "$\rho = 1$ line") where we plot the rank ordered times $t_A$ versus the rank ordered $t_B$; the remaining $1 - |\rho|$ fraction of points were randomly reordered according to the coin method, but maintaining the marginal distributions of $t_A$ and $t_B$. For each target $\rho$, we simulated the joint survival time distribution using the coin method and computed the fraction of points that remained in the "$\rho = 1$ orange line and the Spearman correlation of the simulated data, and plotted them in Fig. S7B. As the result of the previous section anticipated, the fraction of points that remained organized in the $\rho = 1$ line is very approximately equal to the measured Spearman correlation. The discrepancies are due to errors due to sample size. These errors would asymptotically decrease as the number of points increase.

*Survival curves are relatively insensitive to the joint distribution of survival times*
Both the coin method and the window-swap method produce joint distributions of survival times that maintain invarian the marginal distributions and have a given Spearman correlation between -1 and 1. These resulting joint distributions, however, could be quite different, especially as the correlation becomes closer to 1 and -1. Fig. S8A shows the results of the joint distributions for two simulated monotherapies with Hill-curve shaped marginal distributions and Spearman correlation set to 0.25. We can see that there is a remnant of the sorted times in the coin method, delineating a curve that contains 25% of the points, as we showed above. No such line exists for the window swap method. Outside of this line, the remaining points in the coin-method-based joint distribution are independently distributed. In the window-swap-based joint distribution (orange points) all subsets of points are dependent with a Spearman correlation of 0.25. Fig. S8B shows the resulting survival curve corresponing to patients treated with the combination by applying taking the maximum of the survival times under each monotherapy for each simulated patient using the coin simulation (black line) and the window-swap method (orange line). It can be clearly seen that the survival distribution in response to the combination is insensitive to the background joint distribution of survival times.

Fig. S8C and D show similar results as Fig. S8A and B but using the actual monotherapy survival distributions observed in a clinical trial with Erlotinib and Bevacizumab. Fig. S8D shows subtle differences between the coin and window-swap methods, especially for longer survival times. Despite these differences, the two curves are quantitatively very similar, showing that the joint distribution of survival times has at worst a mild effect in the survival distribution in response to the combination treatment resulting from the CDA approach.

#### Fitting the tCDA model to the data

Given the survival probabilities of the monotherapies and their combination, we need to determine how well the tCDA model fits the combination data. For that, we need to find the value of $\rho_s$ in the interval $[-1, 1]$ that results in the best tCDA approximation to the actual combination survival curve. To do so, we can use one of the tCDA implementations (analytical model, coin-based simulation, window swap simulation).

In this work, we deal exclusively with progression-free survival (PFS) curves. But, our methods and approach can be used for any sort of survival metric such as Overall Survival (OS) too.

We estimate the PFS in regular time intervals (we choose 0.05 month intervals) using the scraped raw data via linear interpolation. Next, we find the maximum time point for each trial that we have data for. Then, we compute the minimum of these times across all trials in the combination. We filter all (time, PFS) points with time coordinate values greater than this minimum to ensure that missing values do not interfere with the fitting process.

Now, let us describe fitting using the tCDA analytical model. First, for a given combination, we must assemble progression-free survival tables formatted according to the preceding paragraph. We use Equation 17, and make the simplifying assumption that $\alpha(\rho_s) = \rho_s$. We nominate 200 equally spaced candidate values of $\rho_s$ on the interval $[-1, 1]$ and for each candidate $\rho_s$ we compute its predicted combination survival curve, choosing the $\rho_s$ that minimizes the root mean square error (RMSE) between the observed combination and predicted combination.

As a test that the our fitting procedure captures the right parameter when it should, we simulate survival times for each patient treated with a drug combination using two monotherapies whose survival curves are assumed to follow Hill curves. We then use the coin model to set a correlation between the survival times and simulate the survival in response to the combination as the maximum of the survivals of the monotherapies for each patient. The resulting survival curves for 4 values of correlations are shown as solid blue curves in Fig. S9A. We then fit the best parameter $\alpha$ for the tCDA model from the monotherapies for each correlation shown in the subfigures, and find that the tCDA model fits the simulated survival under the combination treatment extremely well (dashed line in the figure). Furthermore, the fitted values of $\alpha$ resulting from fitting the tCDA model match extremely well the target correlation used in the coin method in the whole range of correlations between -1 and 1 ($R^2$=0.9998) as shown in Fig. S9B. This shouldn't surprise us, as in the previous section we proved that the coin method results in a joint distribution of survival times for which the tCDA model is the exact analytical solution. To show the robustness of the tCDA model to capture the underlying correlation, we followed the same procedure used in Fig. S9 but instead of using the coin method for correlating the joint distribution we used the window swap method. Fig. S10 shows the results. Again the survival distribution for the different imposed correlations (blue solid lines in Fig. S10A is captured very well by the tCDA model with fitted parameter $\alpha$. Fig. S10B shows that the value of $\alpha$ tracks remarkably well the target correlation of the simulations, with an $R^2$=0.991. The results shown in this Figure are remarkable as they show that the analytical tCDA model very closely follow the correlation of the joint distribution even when the joint distribution is not the one underlying the analytical model. The conclusion is that the tCDA model is relatively insensitive to the details of the joint distribution of survival times.

We will now describe fitting using the tCDA simulation methods (either the coin or window swap method). For a given combination, there are two individual monotherapies for which we collect data on. For each of these trials, we have the PFS curve and, therefore, the cumulative distribution (1 - PFS curve is the cumulative distribution). We now sample survival times for hypothetical patients from the cumulative distribution via an inverse transform. The number of hypothetical patients should be the same for each trial in the combination and large (we choose it to be 4 times the number of time points for which we have data for). In practice, it can be set rather arbitrarily. Next, we sort the sampled hypothetical patient PFS times for either monotherapy in decreasing order. Concurrently, we also format the data according to the instructions detailed two paragraphs above. We nominate 200 equally spaced candidate values of $\rho_s$ on the interval $[-1, 1]$. For each candidate $\rho_s$, we shuffle the paired survival times according to the desired simulation method (i.e., window swap or coin) such that the Spearman's correlation between the two vectors of monotherapy PFS times are $\rho_s$. We now have a matrix where each row corresponds to a hypothetical patient and the two columns corresponds to the patients' PFS time under either monotherapy. For each patient, we take the maximum survival time across monotherapies and refer to it as the patients' combination PFS time under the tCDA simulation method. Finally, we can use this distribution of tCDA combination survival times to compute the tCDA estimated PFS survival curve for the candidate $\rho_s$. To compare this estimate to the observed combination PFS curve, we linearly interpolate the tCDA combination estimate on the same time intervals as done for the observed combination. We then compute the RMSE between the tCDA and observed combination and choose the $\rho_s$ that minimizes this error.

The confidence interval for $\rho_s$ in the tCDA model (analytical and simulation models) is computed via a bootstrap approach. We use the empirical inverse transform method to generate new survival times that are distributed according to the original survival curve. The cumulative distribution of the original survival curve is simply 1 minus the original survival curve. We generate resampled data for both of the monotherapies and the combination. To ensure the bootstrapped trial's variability is kept consistent with the original trial, the number of samples taken from the original survival curve is set to be the number of patients in the corresponding clinical trial from where the data was obtained. Then an optimal $\rho$ is computed with the resampled data according to the fitting process of the desired model in a process that is repeated 5000 times to construct the appropriate 95% confidence interval. Specifically, we took the 2.5% and 97.5% quantiles of the empirical distribution for $\rho$'s to be the extremes of the 95% confidence interval.

##### A statistical test to compare the analytical tCDA model to the measured survival probability

Once an optimal $\rho_s$ for the survival curve of the combination is generated under the analytical tCDA model, we must quantitatively determine whether the modeled curve describes the experimentally observed curve sufficiently well. We frame this problem as a classical hypothesis test. The null hypothesis is that there is no significant difference between the analytical tCDA model and the observed combination PFS curves. Combinations that are explainable according to tCDA should fail to reject the null hypothesis. Non-independent drug combinations should reject the null hypothesis.

Before describing the test, we must consider the limits in resolution in the collected data. The resolution of a trial is simply the number of events that can happen which is naturally capped at the number of patients in the trial. There is inherent uncertainty in the observed survival curves for the monotherapies and combined therapies. The resolution of each curve is limited by the number of patients used to construct the curve. The analytical tCDA model uses information from the monotherapy curves, and thus, its resolution is limited by the number of patients in the monotherapy trials. The analytical tCDA resolution is set to be the number of patients in the monotherapy with fewer patients. This is a conservative criterion that reduces the power of the test, but makes the curves that are rejected by the null hypothesis to be more likely to be truly non-independent.

The statistical test is a modified Kolmogorov-Smirnov (KS) test. First, we generate a null distribution of KS statistics. This null distribution quantifies our uncertainty in the collected data and represents the space of possible combination curves that can be realized. We draw $n_{AB}$ samples from the observed combination curve using an inverse transform where $n_{AB}$ is the number of patients in the observed combination. From here, we construct a PFS curve using these re-sampled survival times. Next, we draw $n_{tCDA}$ samples from the observed combination curve using an inverse transform where $n_{tCDA}$ is defined to be the minimum of the number of patients enrolled in either monotherapy. Again, we construct a PFS curve using these re-sampled survival times. Now, we have two re-sampled combination PFS curves. We now compute the KS distance between the two curves. In the event that the two curves do not have the same time coordinates, we use the time coordinates that are closest to each other. In practice, linearly interpolating at pre-specified intervals and this method perform near identically. We then repeat this process of re-sampling and finding the KS distance 100,000 times per combination. This results in a null distribution of KS statistics. When an analytical tCDA survival curve is computed at the $\rho_s$ with minimum RMSE, we draw $n_{tCDA}$ samples from this survival curve via an inverse transform. Then we compute the KS statistic between it and the observed combination survival curve in exactly the same manner that we did when generating the null distribution of KS statistics. The KS distance between the tCDA fitted combination and the true combination is used as a threshold and the p-value is computed by determining the number of samples in the null comparison distribution that have a KS value greater than the threshold.

For a single hypothesis test, we set the p-value threshold at 0.05, a standard level. However, we test multiple hypotheses. We adjust for this by using a Benjamini-Hochberg correction wherein the actual threshold is the individual threshold over the number of trials. This value of 0.0027 is the threshold for rejecting or failing to reject the null model.

#### Code

Code for the tCDA model, bootstrapped confidence intervals, and goodness of fit testing can be found here. The default options for this code is to run the analytical tCDA model, but can also use the simulation tCDA methods to analyze clinical trials.

#### Detailed examination of the tCDA model with Herceptin and chemotherapy combination trial data

We next examine the combination of chemotherapy and Trastuzumab (Herceptin). Chemotherapy consisted of anthracycline plus cyclophosphamide for patients who had never received anthracycline before or paclitaxel for patients who had received adjuvant anthracycline [1]. This therapy is a current treatment for HER-2 overexpressing breast cancers. The Tratuzumab and chemotherapy combination shows a clinical benefit over either individual monotherapy with respect to progression-free survival (PFS) time (Fig. S3A). At around 12 months, the benefit of the combination reduces to that of the Trastuzumab treated group. As shown earlier, the tCDA model predicts the PFS curve $S_{AB}(t)$ of the combination of the two drugs A and B (in this case chemotherapy and Trastuzumab) based on the PFS curves of each of the monotherapies $S_A(t)$ and $S_B(t)$ in the form prescribed by the mathematical expression

$$S_{AB}(t,\rho) = \\ = \begin{cases} S_{\mathsf{max}}(t) + S_{\mathsf{min}}(t)(1 - S_{\mathsf{max}}(t))(1-\rho) & \text{if } 0 \le \rho \le 1 \\ (S_A(t) + S_B(t) - S_A(t)S_B(t))(1 - |\rho|) + \min(1, S_A(t) + S_B(t))|\rho| & \text{if } -1 \le \rho \le 0 \end{cases} \tag{37}$$

where $S_{\mathsf{max}}(t) = \max(S_A(t), S_B(t))$, $S_{\mathsf{min}}(t) = \min(S_A(t), S_B(t))$, and $\rho$ is the correlation over all patients of the times that each patient would have survived if treated with drug A and B independently. As we vary the parameter $\rho$ in Eq.(37) using as $S_A(t)$ and $S_B(t)$ the observed PFS curves of the chemotherapy and Trastuzumab respectively, we span the range of the tCDA predictions forming the cone of possibilities shown in Fig. S3B, where each color corresponds to one value of correlation.

Fig. S3C shows different simulations (using what we called the "Coin method" described earlier in the Supplement) of survival times for different overall correlations in the patient population. Each point in each subpanel of Fig. S3C represents a simulated patient, and the y-axis and x-axis values are the survival times that a given patient would have survived if treated only with Trastuzumab or chemotherapy respectively. The marginal distribution of x-axis and y-axis values in all subfigures follow $S_A(t)$ and $S_B(t)$ curves respectively, and the correlations alluded to in the figure are Spearman's rank correlations. A value of Spearman's correlation equal to 1 traces the "Time Equivalence Curve" between Trastuzumab and chemotherapy while a value of -1 associates the survival times of patients to drugs Trastuzumab and chemotherapy in reverse order. A Spearman's correlation of zero assigns survival times to Trastuzumab and chemotherapy independently of one another. From this joint distribution of patient survival times and by taking the maximum value at each ordered pair, we can construct the empirical estimate of the PFS curve under tCDA model at a given Spearman's correlation. This simulation process is mathematically equivalent to the closed-form expression given in Eq.(37) which we use throughout the paper. We will estimate the tCDA predicted PFS curve of the combination by varying the parameter $\rho$ between $[-1, 1]$ in the analytical model in Eq. (37) and choosing the value of $\rho$ that minimizes the root mean square error (RMSE) between the estimated and observed combination PFS curves.

Returning to the Trastuzumab and chemotherapy trial data, the optimal estimate for the Spearman's correlation is 0.03 and its corresponding 95% confidence interval is $[-0.07, 0.13]$. The confidence intervals are computed via a bootstrap approach where the data is resampled many times to generate an empirical distribution of correlation estimates as described earlier. We can observe that the resulting estimate for the combination PFS curve under tCDA follows the true combination rather well (Fig. S3D). Empirically, using a modified Kolmogorov-Smirnov test statistic described earlier, we fail to reject the null hypothesis that the tCDA model sufficiently describes the observed combination (p-value of 0.46). In this sense, it can be hypothesized that this combination is not inconsistent with the tCDA independent action assumption.

Interestingly, a $\rho = 0$, as we approximately have in this case, reduces Eq. (37) into a mathematical form analogous to that of Bliss independence used in the dose domain, namely $S_{AB}(t, \rho = 0) = S_{\mathsf{max}}(t) + S_{\mathsf{min}}(t)(1 - S_{\mathsf{max}}(t))$. A zero Spearman's correlation indicates that the survival times that can be attributed to each of the treatments are independent of each other in the population. This suggests that in the absence of a stratification strategy to separate metastatic HER-2 overexpressing breast cancer patients into those that respond better to Trastuzumab and those that respond better to chemotherapy, it is better to administer the combination to all such patients rather than trying to assign each patient to the monotherapy that will work best for them. In this scenario, when we are fundamentally unable to decide for a given patient whether Trastuzumab or chemotherapy is better for them and in order to reduce the guesswork in the process, the best strategy is to give the combination for each patient.

#### Additional examples of tCDA model applied to combination trial data

Figures S4 to S6 show several examples of combinations for which the tCDA model yield different Spearman's correlations include 5-FU and Oxaliplatin in advanced pancreatic cancer, Interferon Alfa and Temsirolimus in advanced renal cell carcinoma, Irinotecan Bevacizumab and Panitunumab in advanced colorectal cancer, and other combination chemotherapy regimens. Of course, we must be sufficiently certain that the tCDA model describes well the effect of the combination, and even in that case we must be careful to not over-interpret the meaning of the Spearman's correlation resulting from the tCDA model fit.

#### Dose-space Correlated Drug Action Model

Independent drug action has been mostly explored in the in the context of temporal responses such PFS or OS in clinical trials or pre-clinical research [2, 3]. In this section we investigate the principle of independent drug action at the level of cell cultures and in dose space. To do so we apply the same principles that were applied in previous sections but rather than asking for the survival time of a patient under each of two drugs, we will ask for the survival of a cell to each of two drugs at their respective concentrations and after a fixed time has elapsed since treatment. The assumption in what follows is that each cell will die because of the effect of the most efficacious drug for that cell, and like we did before with the survival time for each patient, we will consider that the doses at which a cell dies with one or another drug are correlated. This correlation could be due to confounding variables. In cell cultures we can think that specific characteristics of a cell such its number of mitochondria [4], its size, its protein content, etc, could be a confounder. We will call this application of correlated response of cells in dose space as Dose Correlated Drug Action, or dCDA.

Like tCDA, dCDA is an independent drug action model for the action of drug combination on cells cultures when studying their dose response curves. There is a body of literature that addresses the problem of predicting the dose-response of a drug combination given the dose responses of the constituent monotherapies under the baseline assumptions of independence or additivity [5]. The most used baseline models are Bliss' independent joint action model [6] and Loewe's dose additivity principle [7]. The latter can be transformed into a quantitative measure called the Combination Index [8], and was the basis of the more general principle of dose equivalence [9]. There are other methods developed for drug combination models such as the Highest Single Agent [10], and some frameworks for their integration into a unified framework [11, 12]. The dCDA starting point is the quantification of killing effect of a therapy on a cancer cell culture using the dose response curve that estimates the survival of a cell culture with respect to a control population. The dCDA model postulates that the effect of two therapies in combination is the same as the effect of the most effective of the two therapies at the single cell level. If we knew a priori which of two therapies yields better results for a given cell in a culture, dCDA postulates that the effect of administering the best monotherapy for that cell would be the same as the effect of administering the combination.

The doses at which cells die vary from cell to cell due to different states in which cells could be, such as abundance of key proteins, number of mitochondria, its size, etc. dCDA assumes that the lethal doses on the same cell in response to each of two drugs could be correlated.

The dCDA framework provides a baseline hypothesis for independent drug action that can be used as a null hypothesis in a statistical test to classify combinations as consistent with dCDA or not consistent with dCDA. Non-dCDA combinations that are of particular interest are antagonistic or synergistic combinations.

#### Notation

For each cell in a cell culture, we will denote by $\delta_A$ and $\delta_B$ the doses at which the cell would be dead at a given time after treatment (typical assays measure viability at a time between 24h to 72h). If treated with drug X (X=A or B) at dose $D_X$, a cell with a lethal dose of $\delta_X$ will be found dead at time T if the effective dose of drug X was larger than $\delta_X$. Under the dCDA model, if a cell with lethal doses $(\delta_A, \delta_B)$ is treated with both drugs at doses $(D_A, D_B)$ then the cell will die if either $\delta_A < D\prime_A$ or $\delta_B < D'_B$. Here $D\prime_A$ is the effective dose of A, enhanced by the fact that the culture is also treated with drug B. Similarly, $D'_B$ is the effective dose of B, enhanced by the fact that the culture is also treated with drug B. We will discuss in a subsequent section how to calculate $D'_A$ and $D'_B$. For a cell to survive, both lethal doses have to be smaller than the equivalent treatment doses.

The dose response curve that a generic cell in a cell culture survives at dose $D_A$, $D_B$ under treatments A, B and combination AB will be denoted by $V_A(D_A)$, $V_B(D_B)$ and $V_{AB}(D_A, D_B)$, and can be formally expressed as

$$V_A(D_A) = \text{Prob}(\delta_A > D_A) \tag{38}$$
$$V_B(D_B) = \text{Prob}(\delta_B > D_B) \tag{39}$$
$$V_{AB}(D_A, D_B) = \text{Prob}(\delta_A > D'_A, \delta_B > D'_B) \tag{40}$$

In practice the probabilities are computed as fractions in a cell culture. For example, the $V_A(D_A)$ is the fraction of cells in culture that are still alive at dose $D_A$ and therefore is the fraction of cells whose lethal dose $\delta_A$ is larger than $D_A$.

The fraction of cells that die is call the inhibited fraction, and is equal to 1 minus the viability, that is:

$$I_A(D_A) = 1 - V_A(D_A) = \text{Prob}(\delta_A < D_A) \tag{41}$$
$$I_B(D_B) = 1 - V_B(D_B) = \text{Prob}(\delta_B < D_B) \tag{42}$$
$$I_{AB}(D_A, D_B) = 1 - V_{AB}(D_A, D_B) = \text{Prob}(\delta_A < D'_A \text{ OR } \delta_B < D'_B) \tag{43}$$
$$\tag{44}$$

The inhibited fractions under drug A or B is equal to the cumulative distribution of the lethal times under those drugs in a cell culture. Therefore, the inhibited fraction or the viability of a cell culture under the effect of drug $X$ can be written in terms of the probability density $\mathcal{P}(\delta_X)$ that a cell has lethal dose $\delta_X$ under treatment $X$ as

$$I_X(D_X) = \int_0^{D_X} \mathcal{P}_X(\delta)d\delta \tag{45}$$

To measure the correlation between lethal doses $\delta_A$ and $\delta_B$ we will use the Spearman's rank correlation. This is computed by ranking the $\delta_A$'s and $\delta_B$'s in a cell culture and computing the correlation between the rankings of $\delta_A$ and $\delta_B$ for the same cell in the cell culture.

#### Formulation of the Dose Correlated Drug Action Model

In this section, we want to derive expressions for the viability of a cell culture under treatment with both drugs A and B at doses $D_A$ and $D_B$ in terms of the viabilities of the monotherapies. To proceed we consider separately the correlated and the uncorrelated cases.

##### Uncorrelated Case

We will consider that the two lethal doses $\delta_A$ and $\delta_B$ for treatments with drugs A and B are uncorrelated because they are statistically independent. (In the general case, the drugs could be uncorrelated but statistically dependent). In this case, the equivalent doses $D'_A$ and $D'_B$ for A and B are equal to the given doses $D_A$ and $D_B$ because the presence of A (resp. B) does not affect A (resp. B) in any way, and Prob($\delta_A \geq D_A$ AND $\delta_B \geq D_B$) = Prob($t_A \geq t$)Prob($t_B \geq t$), which leads to

$$V_{AB}(D_A, D_B, \rho_s = 0) = V_A(D_A)V_B(D_B), \tag{46}$$

Note that in this case, the viability under the two drugs is smaller than the viability under either of the single therapies. If we express the (46) in terms of the Inhibited fraction, we will find the relation $I_{AB}(D_A, D_B, \rho_s = 0) = I_A(D_A) + I_B(D_B) - I_A(D_A)I_B(D_B)$ which has the same expression as (7). The reason why the equation for the survival probability in tCDA is formally the same as the inhibition fraction in dCDA is because the former results from a logical OR condition (patients are alive if $t_A > t$ OR $t_B > t$) whereas in the latter it is the probability of inhibition what follows from an OR condition (cells die if $\delta_A < D_A$ OR $\delta_B < D_B$).

##### Positively correlated case

Let us consider the extreme case in which for any given cell, the lethal dose $\delta_A$ completely determines the lethal dose $\delta_B$, in such a way that the Spearman correlation between the lethal doses under A and B treatments of all cells in a culture is 1. That means that the order of lethal doses of cells under treatment A is the same as the order under treatment B. Take now a cell with lethal doses $\delta_A$ and $\delta_B$. The fraction $V_A(\delta_A)$ of cells that died at dose $D_A = \delta_A$ should be the same as the fraction $V_B(\delta_B)$ of cells that died at dose $\delta_B$ under treatment B. Therefore when the Spearman correlation between lethal doses is $\rho_s = 1$, we have that $V_A(\delta_A) = V_B(\delta_B)$, from where we have the relationship

$$\delta_B = V_B^{-1}(V_A(\delta_A)) := f(\delta_A), \tag{47}$$

$$\delta_A = V_A^{-1}(V_B(\delta_B)) := g(\delta_B) = f^{-1}(\delta_B). \tag{48}$$

Note that the functions $f(\delta_A)$ and $g(\delta_B)$ are increasing functions of $\delta_A$ and $\delta_B$ respectively, because the viabilities $V_A$ and $V_B$ and their inverses $V_B^{-1}$ and $V_A^{-1}$ are decreasing functions of their arguments (the number of surviving cells decreases when we augment the dose of a treatment). Relations (48) express what is known as the dose equivalent model, and was developed [9] in part to account for what is called as sham combination principle [13] which postulates that a combination of a drug with itself should be additive. In our case, the dose equivalence principle appears very naturally in the limiting case of perfectly correlated lethal doses.

It was mentioned earlier that the presence of drug B at dose $D_B$ in the presence of treatment A may enhance the dose of A, $D_A$ in to and effective dose $D'A$. Symmetrically, the effective dose of B will be enhanced by A from $D_B$ to $D'_B$. We will assume that this enhancement is a function of the correlation between doses, in such way that

$$D'_A = D_A + \beta(\rho_s)g(D_B), \tag{49}$$
$$D'_B = D_B + \gamma(\rho_s)f(D_A), \tag{50}$$
$$\tag{51}$$

where $\beta(\rho_s)$ and $\gamma(\rho_s)$ control the enhancement of dose of one drug due to the other drug. These functions are unknown a priori but we only need to know them in the extreme cases of $\rho_s$= 1, 0 and -1. We can find a criterion to find the values of these functions at $\rho_s$=1, by requiring that dCDA be compatible with sham combinations, that is combination of a drug with itself. In fact when we "combine" a drug with itself, drug B *is* drug A, and therefore for any given cell $\delta_B = \delta_A$ and the correlation is 1. As in this case the addition of more of the same drug should result in adding the doses, and both $f(D_A)) = D_A$ and $g(D_B) = D_B$, it follows that sham combination compliance requires that $\beta(1) = 1$ and $\gamma(1) = 1$. The case $\rho_s = 0$ is the other extreme in which there is no effect whatsoever direct or indirect of drug A on drug B, and therefore, the presence of one does not change the effective dose of the other, which suggests that $\beta(0) = 0$ and $\gamma(0) = 0$. The corner case of $\rho_s = -1$ is unusual and it is unclear what the values of these functions should be. In the absence of a clear criterion we will assume $\beta(-1) = 0$ and $\gamma(-1) = 0$.

Coming back to $\rho_s = 1$ we can compute the viability under the combination AB. First we note that we can express Prob($\delta_A \leq D'_A$ AND $\delta_B \leq D'B$) in terms of conditional probabilities as

$$\text{Prob}(\delta_A \geq D'_A \text{ AND } \delta_B \geq D'_B) = \text{Prob}(\delta_A \geq D'_A|\delta_B \geq D'_B)\text{Prob}(\delta_B \geq D'_B) \tag{52}$$
$$= \text{Prob}(\delta_B \geq D'_B|\delta_A \geq D'_A)\text{Prob}(\delta_A \geq D'_A). \tag{53}$$

Suppose now that $V_B(D'_B) < V_A(D'_A)$. This means that at these doses, there are more cancer cells in culture that died with the equivalent dose of drug B than with the equivalent dose of drug A. It also means that the equivalent dose $D'_A$ is smaller than the equivalent dose of $g(D'_B)$, that is $D'_A < g(D'_B)$. Because the order of lethal doses of the cells in culture is the same under both drugs, then all the cells that survived under the effective dose of B, have lethal doses $\delta_A$ larger than $g(D'_B)$. Therefore, $D'_A < g(D'_B) < \delta_B$. It follows that for $V_B(D'_B) < V_A(D'_A)$, Prob($\delta_A \geq D'_A|\delta_B \geq D'_B) = 1$. For $V_A(D'_A) < V_B(D'_B)$ a symmetric argument leads to Prob($\delta_B \geq D'_B|\delta_A \geq D'_A) = 1$. Putting these results together, we have that when there is perfect and positive Spearman correlation, the viability of the combination can be written as

$$V_{AB}(D_A, D_B, \rho_s = 1) = \min(V_A(D'_A), V_B(D'_B)). \tag{54}$$

For an alternative derivation of this result, we can start from the joint probability distribution

$$\mathcal{P}(\delta_A,\delta_B)=\delta(V_A(\delta_A)-V_B(\delta_B))\mathcal{P}_{\mathcal{A}}(\delta_A)\mathcal{P}_{\mathcal{A}}(\delta_A),$$

and compute $V_{AB}$ as

$$V_{AB}(D'_A,D'_B)=\int_{D'_A}^{\infty}\int_{D'_B}^{\infty}\mathcal{P}(\delta_A,\delta_B)d\delta_A d\delta_B \tag{55}$$

$$=\int_{V_A(D'_A)}^{1}\int_{V_A(D'_B)}^{1}\delta(z_A-z_B)dz_A dz_B \tag{56}$$

which leads to Eq. (54). Eq. (54) is known in the literature as the Higher Single Agent model [10], which here results naturally from the fully correlated case of dCDA. However here the arguments of the dose response curves $V_A$ and $V_B$ are the effective doses of the drugs.

When $\delta_A$ and $\delta_B$ are uncorrelated and independent, we have that Prob$(\delta_A \geq D'_A|\delta_B \geq D'_B)$ = Prob$(\delta_A \geq D_A)$ and symmetrically, Prob$(\delta_B \geq D'_B|\delta_A \geq D'_A)$ = Prob$(\delta_B \geq D_B)$, where we used that for $rho_s = 0$ $D'_A = D_A$ and $D'_B = D_B$. In the intermediate case in which we have some correlation $\rho_s$, we will interpolate the conditional probabilities in such a way that the completely correlated and uncorrelated case case match the extreme cases as follows

$$\text{Prob}(\delta_A \geq D'_A|\delta_B \geq D'_B)=\alpha(\rho_s)+(1-\alpha(\rho_s))\text{Prob}(\delta_A \geq D'_A) \text{ if } V_B(D'_B)<V_A(D'_A) \tag{57}$$

$$\text{Prob}(\delta_B \geq D'_B|\delta_A \geq D'_A)=\alpha(\rho_s)+(1-\alpha(\rho_s))\text{Prob}(\delta_B \geq D'_B) \text{ if } V_B(D'_B)\geq V_A(D'_A) \tag{58}$$

where $\alpha(\rho_s = 0) = 0$, $\alpha(\rho_s = 1) = 1$ and the specific functional form in the range $0 \leq \rho_s \leq 1$ will depend on the joint distribution of lethal doses.

Replacing these results in (52) and (53) and using (40) we find that the viability under the combination AB in the case of $0 \leq \rho_s \leq 1$ for the dCDA model is

$$V_{AB}(D_A,D_B,0\leq\rho_s\leq 1)=V_A(D_A)V_B(D_B)(1-\alpha(\rho_s)) + \begin{cases} V_A(D_A+g(D_B))\alpha(\rho_s) & \text{if } V_A(D_A+g(D_B))<V_B(D_B+f(D_A)) \\ V_B(D_B+f(D_A))\alpha(\rho_s) & \text{if } V_A(D_A+g(D_B))\geq V_B(D_B+f(D_A)). \end{cases} \tag{59}$$

###### Negatively correlated case

Let us consider the other extreme case in which for any given cell the lethal dose $\delta_A$ completely determines the lethal dose $\delta_B$, in such a way that the Spearman correlation between lethal doses under A and B treatments of all all cells is -1. That means that the order of lethal doses of cells under treatment A is opposite to the order under treatment B. Take now a cell with lethal doses $\delta_A$ and $\delta_B$. The fraction $V_A(\delta_A)$ of cells that have higher lethal doses than $\delta_A$, should be the same as the fraction $1 - V_B(\delta_B)$ of cells that had smaller lethal doses than $\delta_B$. Therefore when the Spearman correlation between lethal doses is $\rho_s = -1$, we have that (see Fig.)

$$\delta_B = V_B^{-1}(1-V_A(\delta_A)) := h(\delta_A). \tag{60}$$

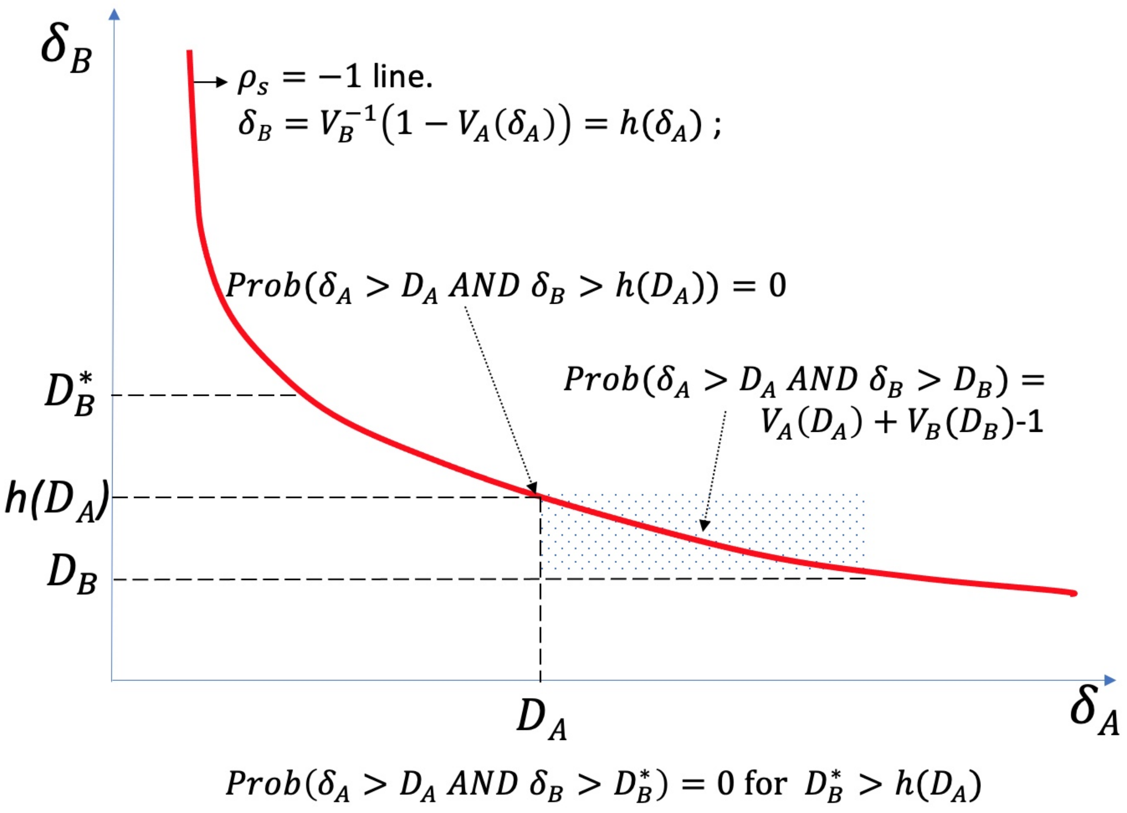


**Figure S2: Case $\rho_s = -1$ for dCDA.**

In this extreme case we can compute the survival time under the combination AB. Recall that we assumed that in this case $D'_A = D_A$ and $D'_B = D_B$. First we note that there for every dose $D_A$ of drug A, there is a dose $h(D_A)$ of drug B such that none the cells that lethal doses larger than $D_A$ (and therefore survived drug A), have lethal doses higher than $h(D_A)$ (and therefore survived B). Therefore for $V_{AB}(D_A, h(D_A)) = 0$ Figure S2 shows that for $D_B < h(D_A)$, the fraction of cells that survived doses $D_A$ and $D_B$ under A or B is equal to $V_A(D_A) + V_B(D_B) - 1$. Figure S2 also shows that for $D_B > h(D_A)$, no cell can survive both $D_A$ and $D_B$. Therefore for $\rho_s = -1$, we have that

$$V_{AB}(D_A, D_B, \rho_s = -1) = \begin{cases} 0 & \text{if } V_A(D_A) + V_B(D_B) < 1 \\ V_A(D_A) + V_B(D_B) - 1 & \text{if } V_A(D_A) + V_B(D_B) \geq 1 \end{cases} \quad (61)$$

$$= \max(0, V_A(D_A) + V_B(D_B) - 1). \quad (62)$$

In the range of negative correlations between -1 and 0, we can interpolate between the cases of $\rho_s = 0$ and $\rho_s = -1$, which yields

$$V_{AB}(D_A, D_B, -1 \leq \rho_s \leq 0) = V_A(D_A)V_B(D_B)(1 - \alpha(\rho_s)) + \alpha(\rho_s)\max(0, V_A(D_A) + V_B(D_B) - 1) \quad (63)$$

where $\alpha(\rho_s = 0) = 0$, $\alpha(\rho_s = -1) = 1$, and the specific dependence of $\alpha(\rho_s)$ on $\rho_s$ depends in the joint distributions of times $\delta_A$ and $\delta_B$.

##### Integrated expression for the viability of cells under the dCDA model

The formulas of the previous subsection can be integrated in one simple integrated formula. The expression for the viability under the dCDA model can be written as:

$$\begin{aligned} V_{AB}(D_A, D_B, \rho_s) &= (1-\alpha(\rho_s))V_A(D_A)V_B(D_B) \\ &+ \alpha(\rho_s)\begin{cases} \min(V_A(D_A+g(D_B)), V_B(D_B+f(D_A))) & \text{if } 0 \le \rho_s \le 1 \\ \max(0, V_A(D_A)+V_B(D_B)-1) & \text{if } -1 \le \rho_s \le 0 \end{cases} \end{aligned} \tag{64}$$

The model is not complete until we specify the function $\alpha(\rho_s)$, which is unknown except in the border cases $\rho_s$ =-1, 0, 1 for which $\alpha(\rho_s)$= is 1, 0, 1. As an approximation, we can assume that $\alpha(\rho_s) = |\rho_s|$. In the next section we will show that this is indeed the correct function for a particular joint distribution for $\delta_A$ and $\delta_B$.

##### There is a joint distribution of lethal doses for which $\alpha(\rho_s) = |\rho_s|$

There are many possible joint distributions of $\delta_A$ and $\delta_B$ that yield the same Spearman correlation, and for each such distribution there may be a different function $\alpha(\rho_s)$. Therefore from the perspective of defining uniquely the dCDA model, it is necessary to specify the joint distribution of lethal doses. In this section we will choose such a distribution and show that for it, $\alpha(\rho_s) = |\rho_s|$, which we will call the dCDA joint lethal doses probability density, or dCDA distribution for short.

This distribution can be thought of as a mixture distribution where a fraction $0 \le \alpha \le 1$ of ($\delta_A$, $\delta_B$) pairs are drawn from the line $\delta_B = f(\delta_A)$ for $0 \le \rho_s \le 1$ or the line $\delta_B = h(\delta_A)$ for $1- \le \rho_s \le 0$, and the other $1-\alpha$ fraction are independently drawn from the distributions $\mathcal{P}_A(\delta_A)$ and $\mathcal{P}_B(\delta_B)$. Formally this can be written as

$$\begin{aligned} \mathcal{P}(\delta_A, \delta_B) &= (1-\alpha)\mathcal{P}_A(\delta_A)\mathcal{P}_B(\delta_B) \\ &+ \alpha\begin{cases} \delta(V_A(\delta_A) - V_B(\delta_B))\mathcal{P}_A(\delta_A)\mathcal{P}_B(\delta_B) & \text{if } 0 \le \rho_s \le 1 \\ \delta(V_A(\delta_A) + V_B(\delta_B) - 1)\mathcal{P}_A(\delta_A)\mathcal{P}_B(\delta_B) & \text{if } -1 \le \rho_s \le 0 \end{cases} \end{aligned} \tag{65}$$

Following the same derivation as in the section "There is a joint survival times distribution for which $\alpha(\rho_s) = |\rho_s|$" it can be shown that for the distribution given in (65), the dCDA model is exact and corresponds to $\alpha(\rho_s) = |\rho_s|$. Therefore we will use this specification of $\alpha(\rho_s)$ in the interpolation between the cases $\rho_s$ =-1, 0 and 1 for the dCDA model.

##### The dCDA model

We saw in an earlier section that the dCDA framework accommodates a spectrum of models that goes from the Bliss independence model at $\rho_s = 0$ to the Higher Single Agent model at $\rho_s = 1$ modified to accommodates for the case of sham combinations. We also noted that at $\rho_s = 1$ the Dose Equivalence Principle appears naturally as the relation between the lethal doses. The model is sham combination compliant because when the two drugs A and B are the same, $\rho_s = 1$, and in this case the model reduces $V_{AB}(D_A, D_B) = V_A(D_A + D_B)$ as required. For the specification of a joint distribution of lethal doses given in Eq.(65) we have that $\alpha(\rho_s) = |\rho_s|$. Therefore we can write the final dCDA model to be used in the paper as:

$$\begin{aligned} V_{AB}(D_A, D_B, \rho_s) = (1 - |\rho_s|)V_B(D_B)V_A(D_A) \\ + |\rho_s| \begin{cases} \min(V_A(D_A + g(D_B)), V_B(D_B + f(D_A))) & \text{if } 0 \leq \rho_s \leq 1 \\ \max(0, V_A(D_A) + V_B(D_B) - 1) & \text{if } -1 \leq \rho_s \leq 0 \end{cases} \end{aligned} \tag{66}$$

#### Parameterization of the viabilities using Hill Curves

We can parameterize the dose-viability curve for a generic drug A ($V_A(D_A)$) by a Hill curve with free parameters $n_A$ and $k_A$. The generic Hill curve equation often used in modeling dose response curves for drug X (X = A or B) at dose $D_X$ is $V_X(D_X) = \frac{1}{1+(\frac{D_X}{k_X})^{n_X}}$ where $k_X$ is dose at 50% effect and $n_X$ controls the steepness of the curve.

To estimate $k_A$, we use the experimental doses and their respective viabilities and linearly interpolate between the experimental values to obtain more data points, then regress viabilities on doses to arrive at the estimated slope and intercept. The slope and intercept of this line, $s_A$ and $i_A$ are used to compute the EC50 or $k_A$ value. Note that for ease of computation viabilities are transformed from 0-1 to 0-100. Therefore, we estimate $k_A$ by $\frac{50-i_A}{s_A}$. Then, using the functional form of the Hill curve given by the expression above for $V_A(D_A)$, we numerically estimate $n_A$ (See Code). We slowly increase the guessed value of $n_A$, compute the resulting Hill curve and find the root mean square error (RMSE) at the points for which we have experimental data. The $n_A$ value with lowest RMSE is chosen. The same procedure can be done for $k_B$ and $n_B$. Under these parameterizations, some of the functions that we described above can be explicitly written. For example the functions for equivalent doses $f(D_A)$ and $g(D_B)$ are

$$f(D_A) = k_B(\frac{D_A}{k_A})^{\frac{n_A}{n_B}} \tag{67}$$

$$g(D_B) = k_A(\frac{D_B}{k_B})^{\frac{n_B}{n_A}} \tag{68}$$

#### Fitting the dCDA model to the data

The optimal estimate for the observed combination viability under the dCDA model can be found by estimating the parameter $\rho_s$, which is constrained between $[-1, 1]$, that best fits the experimental data. We first estimate the Hill curve parameters for each of the two individual drugs as described in the section above. Next, we nominate 200 equally spaced candidate values of $\rho_s$ on the interval $[-1, 1]$ and for each candidate $\rho_s$ we compute its predicted combination viability curve using Equation 66. From these 200 possible combination viability curves, we choose the $\rho_s$ corresponding to the curve that minimizes the RMSE between the esitmated and observed combination viabilities.

Now, we have an initial estimate for $\rho_s$ for a given combination using all the data points, but we have yet to account for possible outliers in the experimental data. We must find and remove outliers from the data and then re-estimate $\rho_s$. We first perform a linear regression between the observed viabilities and the initial combination estimates. Then, we computed the externally studentized residuals which should come from a t-distribution with $n-3$ (because only 1 covariate) degrees of freedom where $n$ is the number of points in the data set. Outliers were defined as points with absolute value greater than a predetermined cutoff $(1-\frac{0.05}{2n})$ based on $n$, the number of total points. There was less than 1 outlier on average found per combination (0.615 outliers/combination). Once we remove the outliers from the data, we estimate $\rho_s$ using the exact same procedure as in the preceding paragraph. The correlation estimate did not change for 84% of the combinations after removing outliers which suggests that the dCDA model is robust. The complete data regarding number of outliers, and before and after removing outliers optimal correlation estimate can be found in Supplemental Table 2. We use the outlier removed correlation estimates for further analysis.

#### Quantification of combinations not accounted for by the dCDA model

Excess over Bliss (EOB) is a commonly used metric for assessing synergistic and antagonistic drug combination effects. It is calculated as $EOB = V_A(D_A)V_B(D_B) - V_{AB}(D_A, D_B)$. By computing EOB, the assumption is that Bliss independence represents the null model. We tested the assumption that Bliss independence is a valid null model by comparing the observed combination viabilities to the Bliss independence viabilities ($\rho_s = 0$ in the dCDA model) and computing a p-value based on a two-sample paired t-test. A p-value $> 0.01$ suggests that the EOB condition adequately provides a measure of synergy/antagonism in the given combination.

We introduce a more general form of deviation from the dCDA null model: the Excess over CDA (EOCDA) - which, for different values of $\rho_s$, ranges from Bliss independence to the sham compliant HSA model depending on the optimal $\rho_s$ value. EOCDA is measured by the difference between the estimated combination viabilities under dCDA and the observed combination viability. A non-zero EOCDA indicates some sort of interaction between the drugs: EOCDA greater than 0 suggests synergy and an EOCDA less than 0 suggests antagonism. The magnitude of the difference indicates the deviation from independent drug action.

Once an optimal estimate for the combination viabilities has been found under dCDA, goodness of fit must be assessed. A two-sample paired t-test was used to compute a p-value indicating evidence against the null hypothesis that the CDA estimate describes the observed combination. For a single hypothesis test, we set the p-value threshold at 0.05, a standard level. However, we test multiple hypotheses. We adjust for this by using a Benjamini-Hochberg correction wherein the actual threshold is the individual threshold over the number of trials. This value of 0.0019 is the threshold for rejecting or failing to reject the null model. A p-value less than this threshold suggests that the null hypothesis should be rejected and that there exists a non-dCDA process that better describes the combination. Results for the dCDA model on the tested combinations can be found in Supplemental Table 2.

#### Identifying locally possibly synergistic or antagonistic dosages in a globally independent joint action drug combinations

We want to identify which, if any, points in a given dCDA independent-joint-action drug combination are possibly acting non-independently (i.e., possibly acting synergistically or antagonistically). Here, a point refers to a given dose of each monotherapy and its associated optimal dCDA estimated viability and the observed viability. We can plot the estimated and observed viability (Fig 3D, F) and visually, those points that are far from the identity are candidates to be locally synergistic or antagonistic dosages. We will formalize this notion now.

First, let us compute the residuals across all points (estimate - observed). We then assume that this distribution is asymptotically normal and estimate the mean ($\hat{\mu}$) and variance ($\hat{\sigma^2}$) of its corresponding normal distribution using the MLE estimate for these parameters.

We remove a given point $i$ from the set. Then, we re-estimate the optimal $\rho$ using the dCDA model and arrive at a new optimal estimate for the viabilities. Next, using the newly optimal $\rho$, we compute the estimated value at the held out point and the residual at this point ($r_i$). Finally, we compute the z-score of this point ($\frac{\hat{\mu}-r_i}{\hat{\sigma}}$) where $\hat{\sigma}$ is the square root of $\hat{\sigma^2}$. Then, we compute the two-sided p-value using this z-score in reference to a standard normal distribution. If the p-value is greater than 0.01, then we fail to reject the null hypothesis that point $i$ is consistent with the dCDA model.

##### Code

Code for the dCDA model can be found here.

#### Supplementary Tables

Table found as attached supplementary Excel file.
**Table S1: tCDA Model Results.** *Sheet 1:* Clinical trial results with Trial ID's and corresponding results for each clinical trial combination.

Table found as attached supplementary Excel file.
**Table S2: dCDA Model Results.** Experimentall tested cell line combinations results with experiment IDs.

Table found as attached supplementary Excel file.
**Table S3: Clinical Trial Data Sources.** Guide describing the sources from which the clinical trial data was assembled.

File found as attached supplementary Excel file.
**File S1: Clinical Trial PFS Data.** Aggregated raw clinical trial progression-free survival data.

File found as attached supplementary Excel file.
**File S2: Cell Line Combination Experiment Data.** Raw data for the experimentally tested combinations.

#### Supplemental Figures

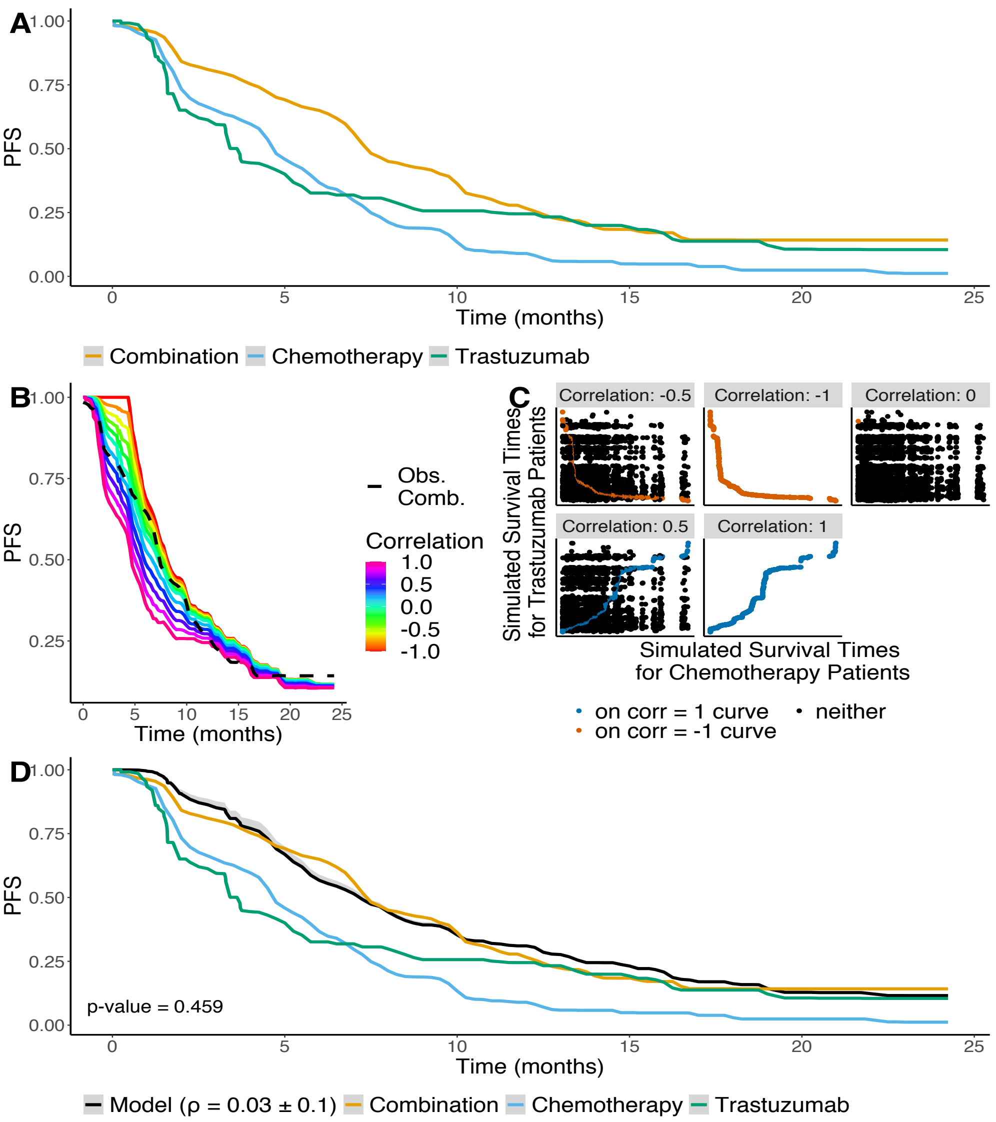


**Figure S3:** Correlated Drug Action explains the benefit of Trastuzumab and Chemotherapy in Metastatic HER-2 Overexpressing Breast Cancer. **A)** Progression Free Survival (PFS) as a function of time for either individual therapy option (Trastuzumab - green, Chemotherapy - blue) and their combination (yellow) in patients with HER-2 overexpressing breast cancer. **B)** Range of possible survival curves for the combination under tCDA. The observed combination (black) falls within the lines of the predicted field. **C)** For a given Spearman's correlation, each point represents a pair of possible PFS times associated with each simulated patient. The parallel maximum, maximum survival time between the coordinates of each point, of this paired vector of PFS times defines the combination PFS curve shown in **B**. The absolute value of the Spearman's correlation corresponds to the fraction of points that lie on the correlation -1 or 1 curves (Fig. S7). **D)** Estimate of the combination under the tCDA model (black) and in grey are the PFS curves for the 95% confidence interval of $\rho_s$ ($0.03 \pm 0.1$, p-value $=$ 0.459). This suggests that the tCDA model sufficiently describes the effect of the combination.

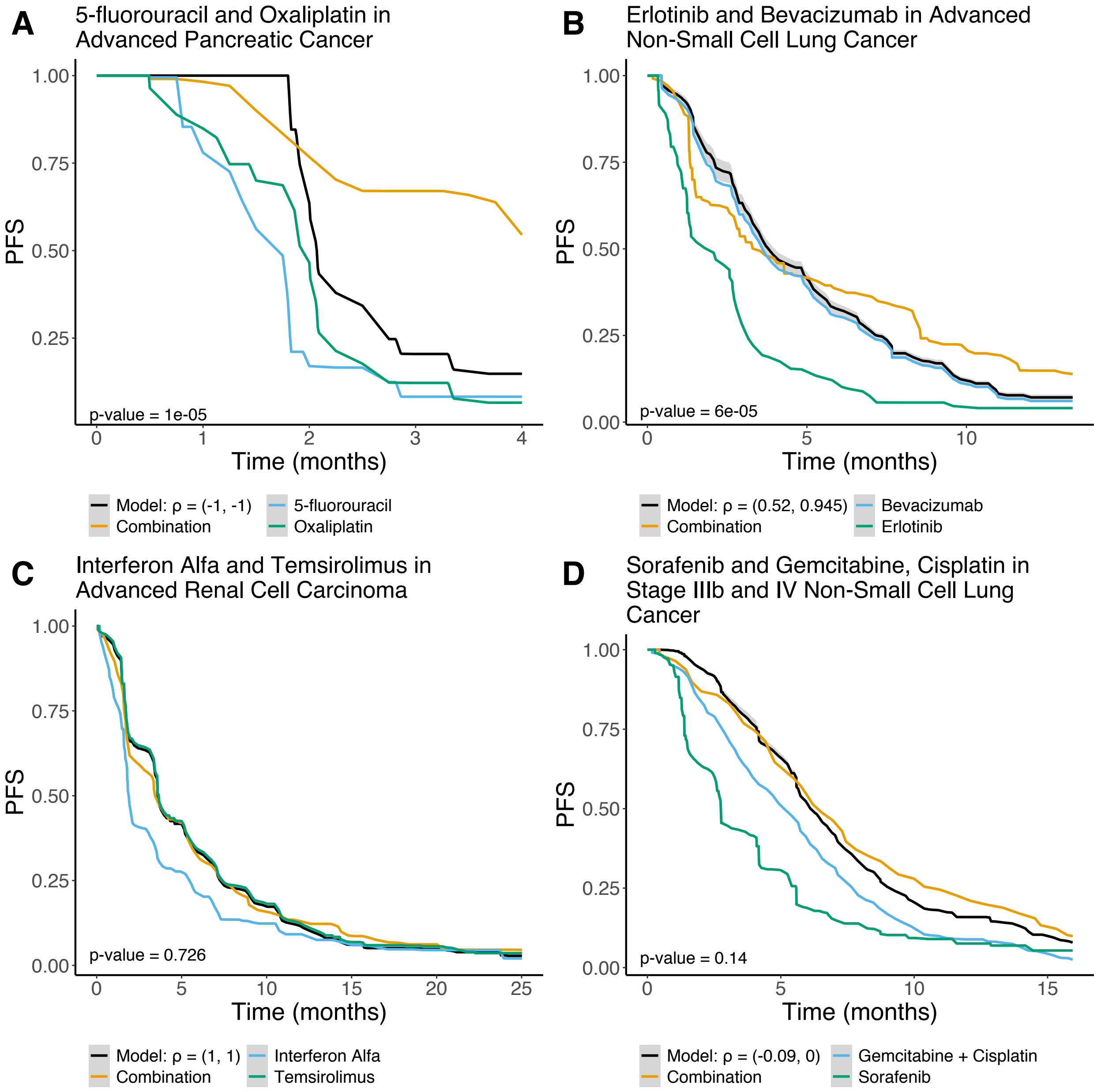


**Figure S4: tCDA model results. A)** Combination of 5-fluorouracil and Oxaliplatin in advanced pancreatic cancer. **B)** Combination of Erlotinib and Bevacizumab in advanced non-small cell lung cancer. **C)** Combination of Interferon Alfa and Temsirolimus in advanced renal cell carcinoma. Since the optimal Spearman's correlation is 1, the tCDA converges to simply following the monotherapy PFS curve with higher PFS at each time step. The model estimate (black) is equivalent to that of Temsirolimus monotherapy (green), the better performing monotherapy. **D)** Combination of Sorafenib and Gemcitabine, Cisplatin in Stage IIIb and IV Non-Small Cell Lung Cancer. **A, B, C, D)** For a given clinical trial, the estimated combination survival curve (black), associated 95% confidence interval (grey) under tCDA, p-value regarding goodness-of-fit, and optimal Spearman's correlation estimate are shown alongside the individual monotherapies and observed combination PFS curves.

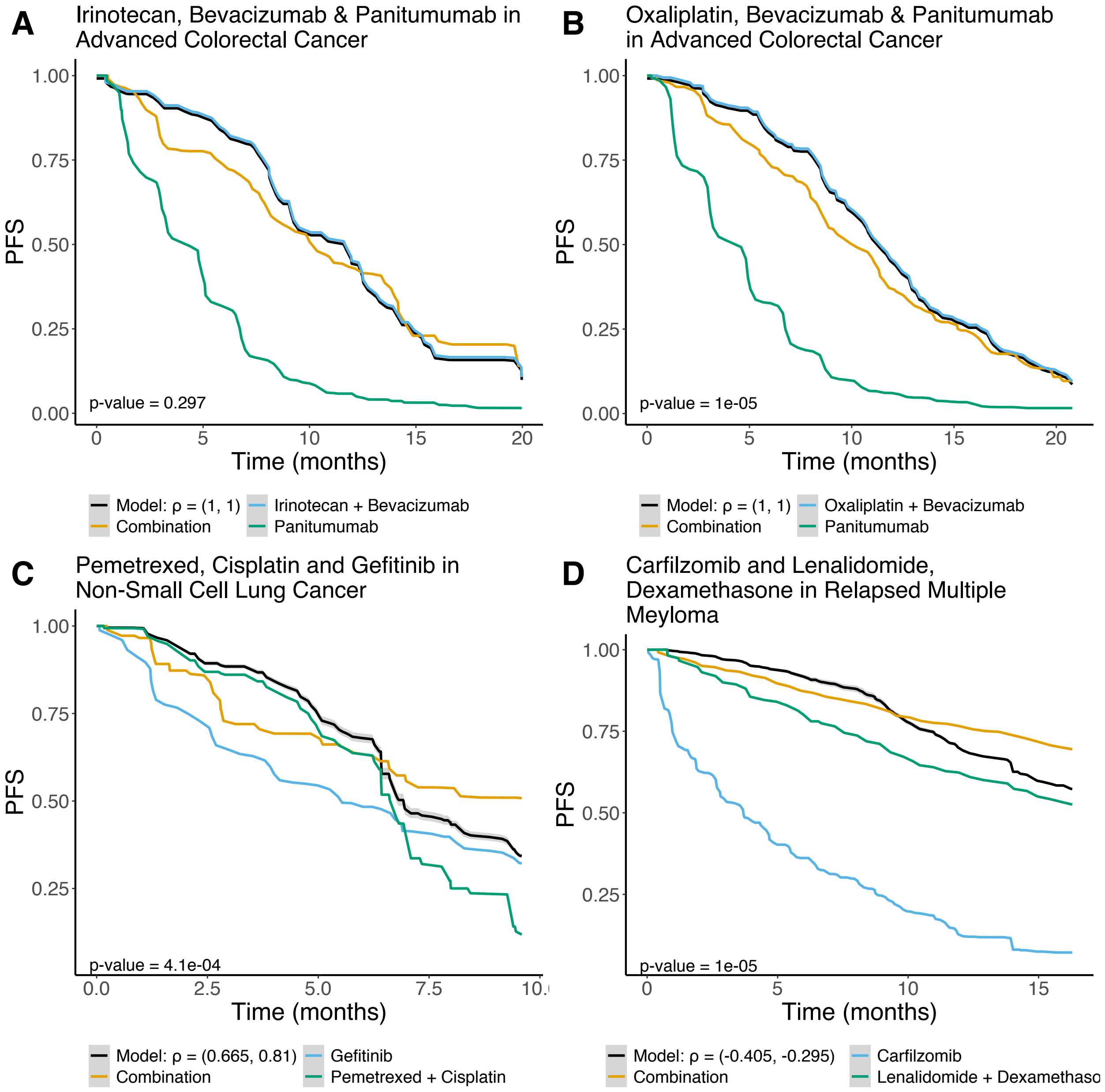


**Figure S5: tCDA model results.** **A)** Combination of Irinotecan, Bevacizumab, and Panitumumab in advanced colorectal cancer. **B)** Combination of Oxaliplatin, Bevacizumab and Pantiumumab in advanced colorectal cancer. **A,B)** Since the optimal Spearman's correlation is 1, the tCDA converges to simply following the monotherapy PFS curve with higher PFS at each time step. Therefore, the model estimate (black) is equivalent to that of the better performing monotherapy (blue). **C)** Combination of Pemetrexed, Cisplatin, and Gefitinib in non-small cell lung cancer. **D)** Combination of Carfilzomib, Lenalidomide and Dexamethasone in relapsed multiple myeloma. **A, B, C, D)** For a given clinical trial, the estimated combination survival curve (black), associated 95% confidence interval (grey) under tCDA, p-value regarding goodness-of-fit, and optimal Spearman's correlation estimate are shown alongside the individual monotherapies and observed combination PFS curves.

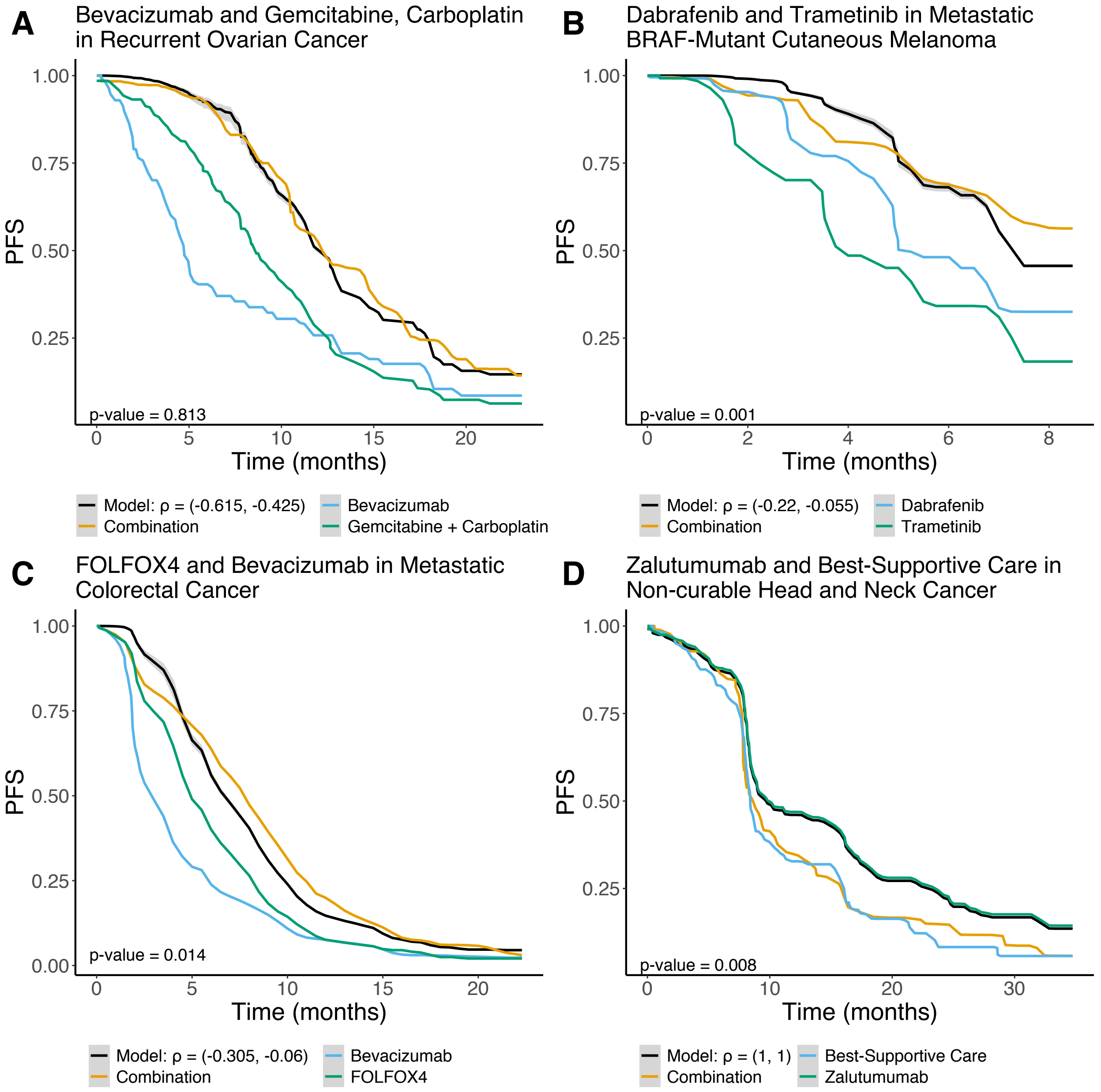


**Figure S6: tCDA model results. A)** Combination of Bevacizumab, Gemcitabine, and Carboplatin in recurrent ovarian cancer. **B)**Combination of Dabrafenib and Trametinib in metastatic BRAF-mutant cutaneous melanoma. **C)** Combination of FOLFOX4 and Bevacizumab in metastatic colorectal cancer. **D)** Combination of Zalutumumab and best-supportive care in non-curable head and neck cancer. Since the optimal Spearman's correlation is 1, the tCDA converges to simply following the monotherapy PFS curve with higher PFS at each time step. The model estimate (black) is equivalent to that of the Zalatumumab PFS curve (green), the better performing monotherapy. **A, B, C, D)** For a given clinical trial, the estimated combination survival curve (black), associated 95% confidence interval (grey) under tCDA, p-value regarding goodness-of-fit, and optimal Spearman's correlation estimate are shown alongside the individual monotherapies and observed combination PFS curves.

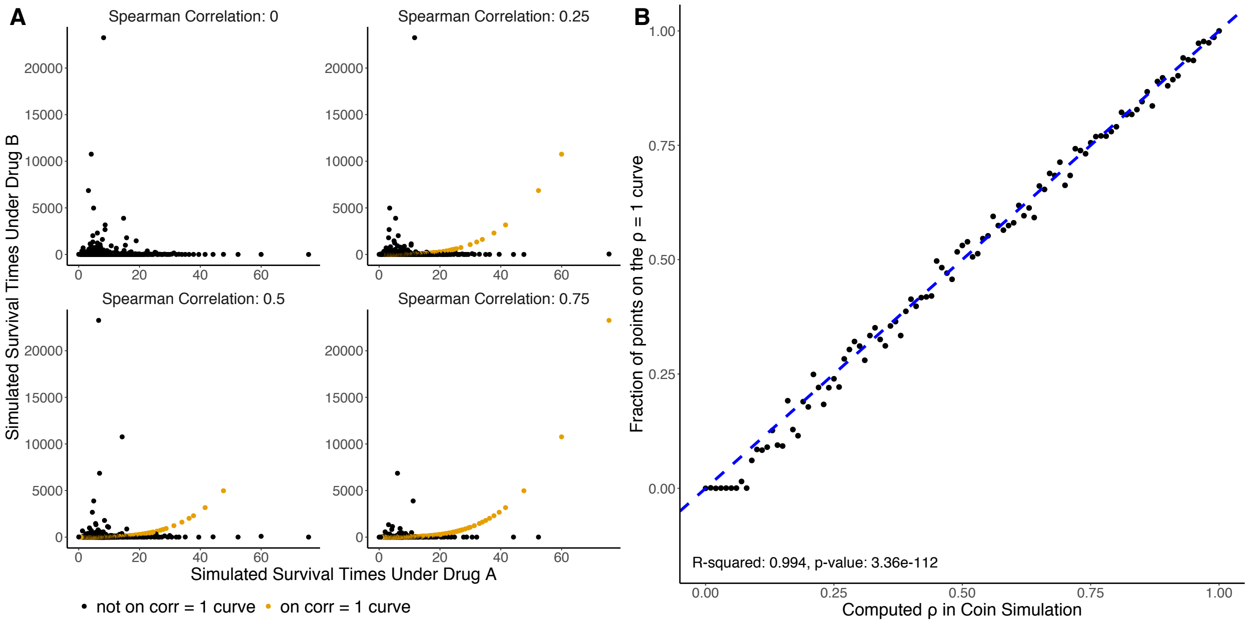


**Figure S7: Rank correlation under the coin method of simulation describes the fraction of points that lie on the perfect rank correlated curve. A)** Simulated data was created and the data was randomized with four distinct values of Spearman's correlation under the coin method of simulation (See Methods). The points that lie on the correlation 1 curve are colored in orange. As the correlation increases, so too does the number of points that lie on the Spearman correlation 1 curve. **B)** An alternative interpretation of the Spearman correlation in the coin method is that it represents the fraction of points that lie on the Spearman correlation equals one curve (R-squared = 0.994 , p-value = 3.4e-112). The dashed blue line is the identity.

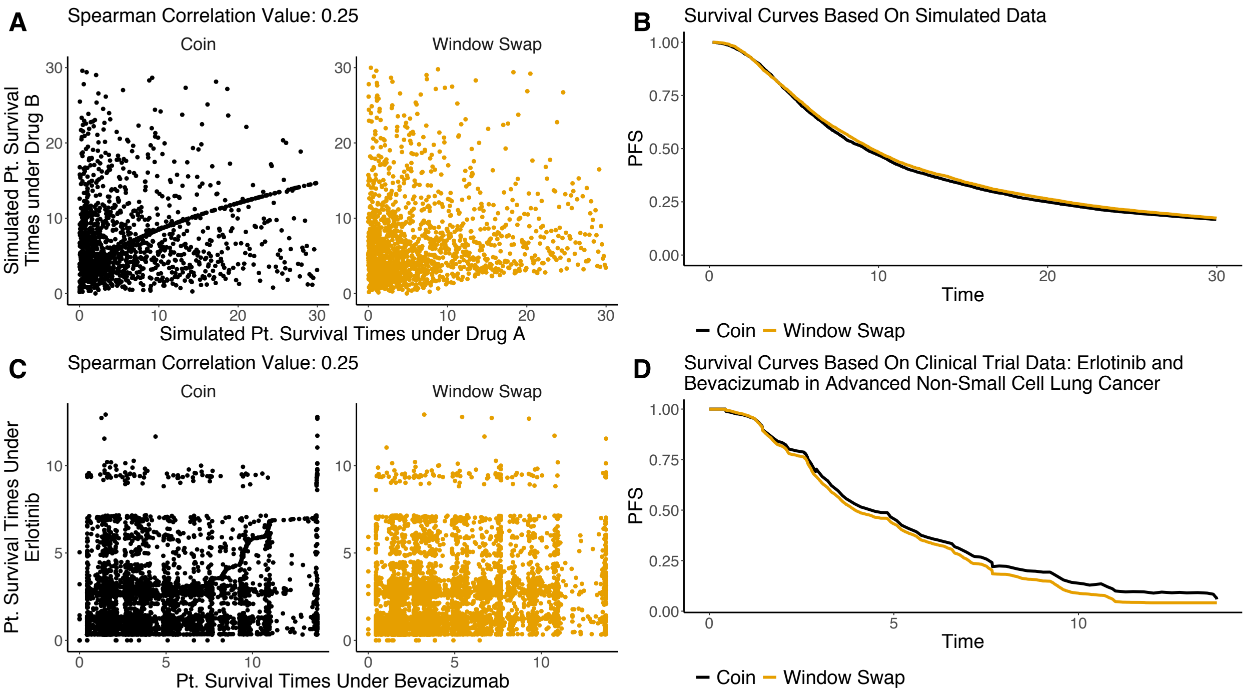


**Figure S8: Differences in performance of simulation methods under synthetic and real data. A)** Simulated data was created to create survival times under drug A and drug B. The times were randomized with Spearman correlation 0.25 under two different simulation methods - coin and window swap (see Methods) . For each simulated patient, their respective times of survival under the individual therapies are shown. **B)** The corresponding survival curves produced by both simulation methods (panel **A**) are shown and are highly concordant despite clear differences between the joint distributions shown in **A**. **C)** Data of the individual therapies from the clinical trial of Erlotinib and Bevacizumab in Advanced Non-Small Cell Lung Cancer were taken and randomized with Spearman correlation 0.25 under the coin and window swap simulation methods. The resulting survival times for a patient are represented as points in the plots. **D)** The resulting survival curves using the data from **C** are shown. The noise inherent within real data contributes to the difference between the survival curves.

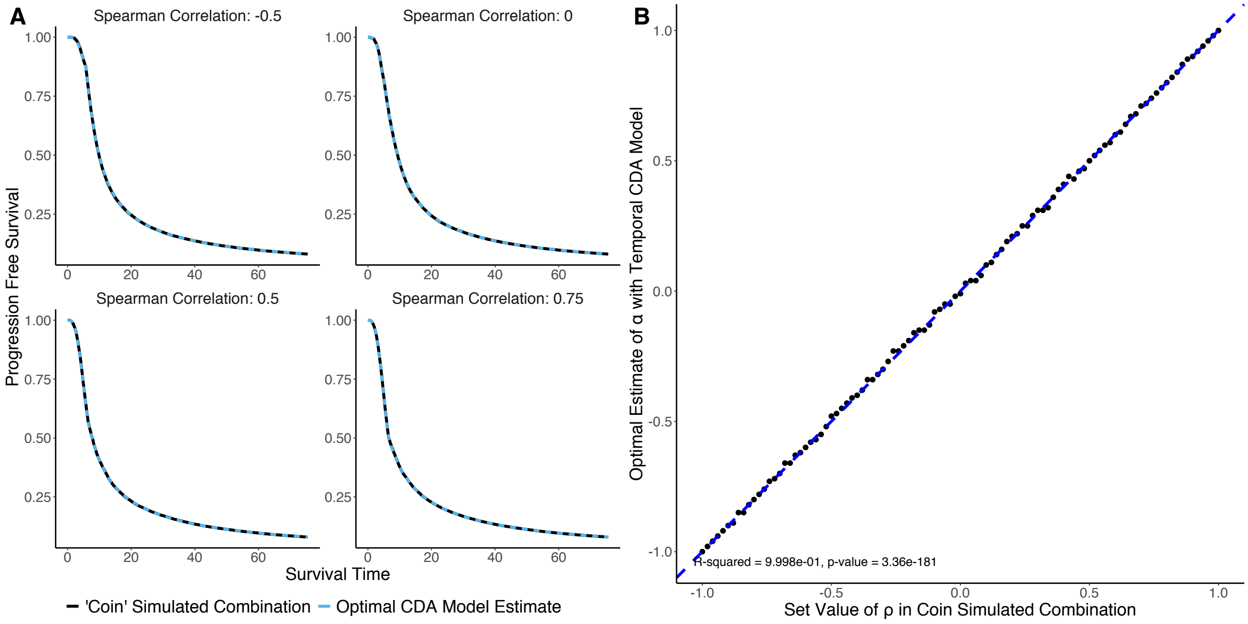


**Figure S9: tCDA model is equivalent to the coin method of simulation. A)** A drug combination was simulated using the coin method of randomization with specified Spearman's correlation and the tCDA model was employed to find a best estimate. The coin simulated combination and tCDA model estimates are highly concordant. **B)** Combination results were simulated with the coin method (See Methods) and estimated with the tCDA model. This was done for Spearman correlations between -1 and 1 with a step-size of 0.02. The input Spearman's correlation for the coin method and the output optimal estimate for the free parameter $\alpha$ in the tCDA model was plotted (R-squared = 0.9997, p-value = 4.36e-178). The dashed blue line is the identity.

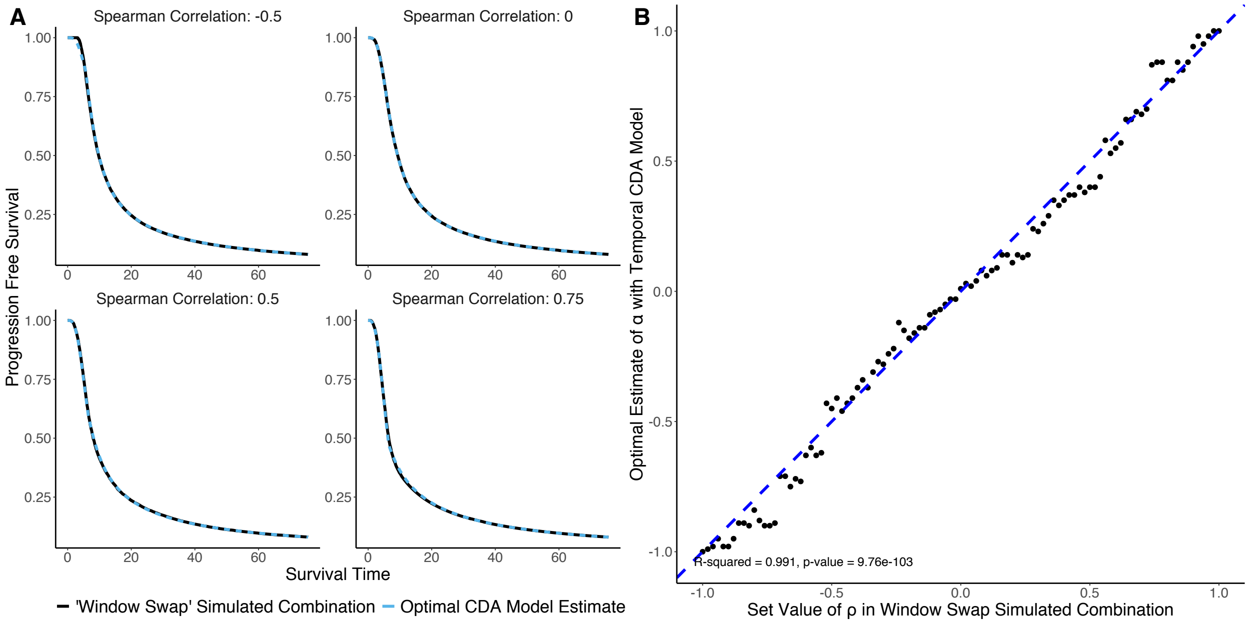


**Figure S10: tCDA model captures much of the variance produced by the window swap method of simulation**. **A)** A drug combination was simulated using the window swap method of randomization with specified Spearman's correlation and the tCDA model was employed to find a best estimate. **B)** Combination results were simulated with the window swap method (See Methods) and estimated with the tCDA model. This was done for Spearman correlations between -1 and 1 with a step-size of 0.02. The input Spearman's correlation for the window swap method and output optimal estimate for the free parameter $\alpha$ in the tCDA model were plotted (R-squared = 0.992, p-value = 8.69e-106). The dashed blue line is the identity.

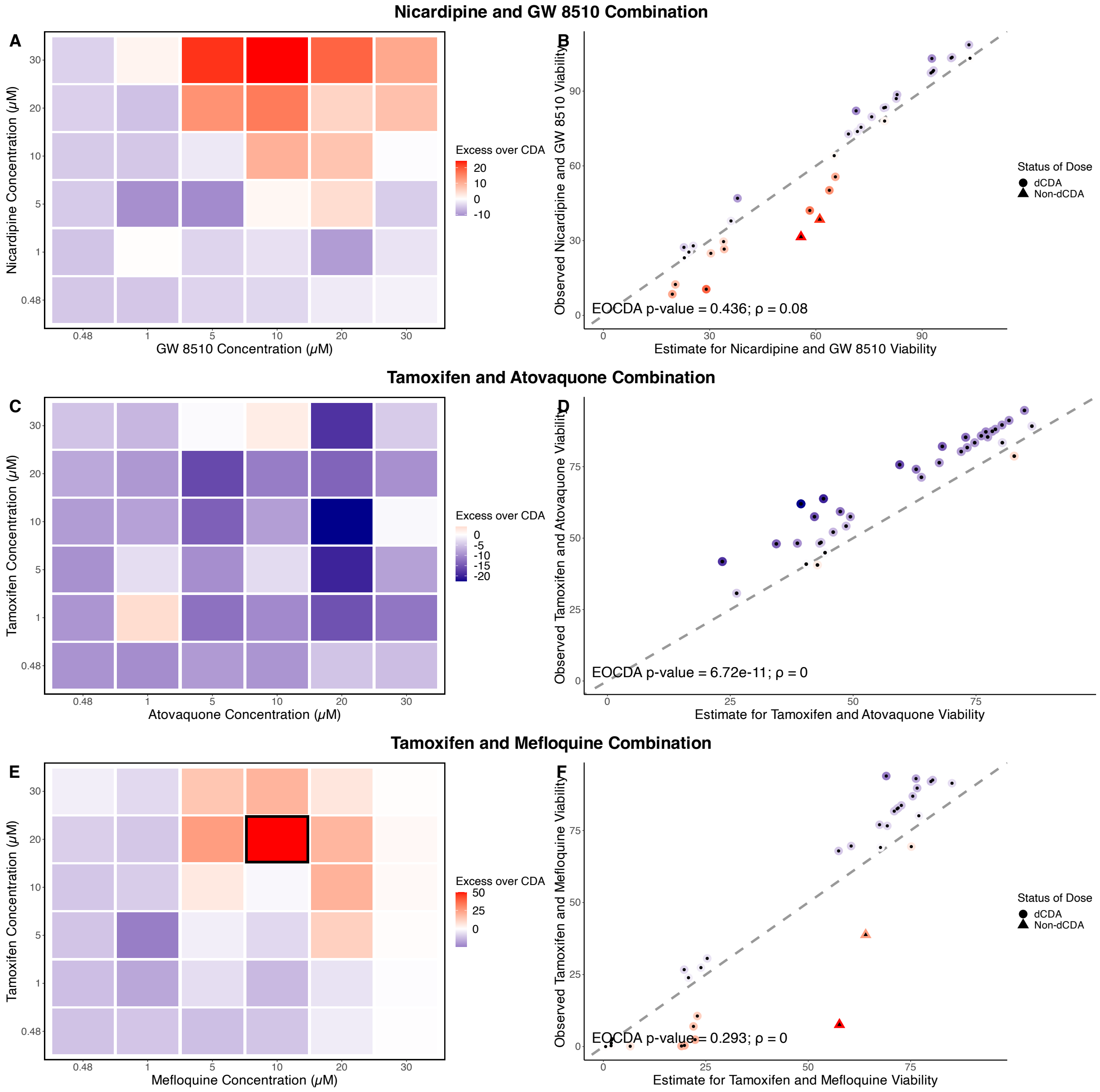


**Figure S11: dCDA model results.** Each row corresponds to a given combination. **A, B)** Nicardipine and GW 8510 combination in MCF7 cells collected after 24 hours. **C, D)** Tamoxifen and Atovaquone combination in MCF7 cells collected after 24 hours. **E, F)** Tamoxifen and Mefloquine combination in MCF7 cells collected after 24 hours. **A,C,E)** Heatmap of excess over CDA is shown with outlier cells bordered in black. **B, D, F)** Comparison of combination estimates and observed viabilities along with goodness-of-fit (GoF) p-value and corresponding optimal Spearman correlation's estimate. Points are colored with the same scale as its corresponding EOCDA matrix. If the GoF p-value > 0.01 for the overall combination, then each point (i.e., dose) is classified as following the dCDA model or not (i.e., likely synergistic or antagonistic behavior) (See Supplement).

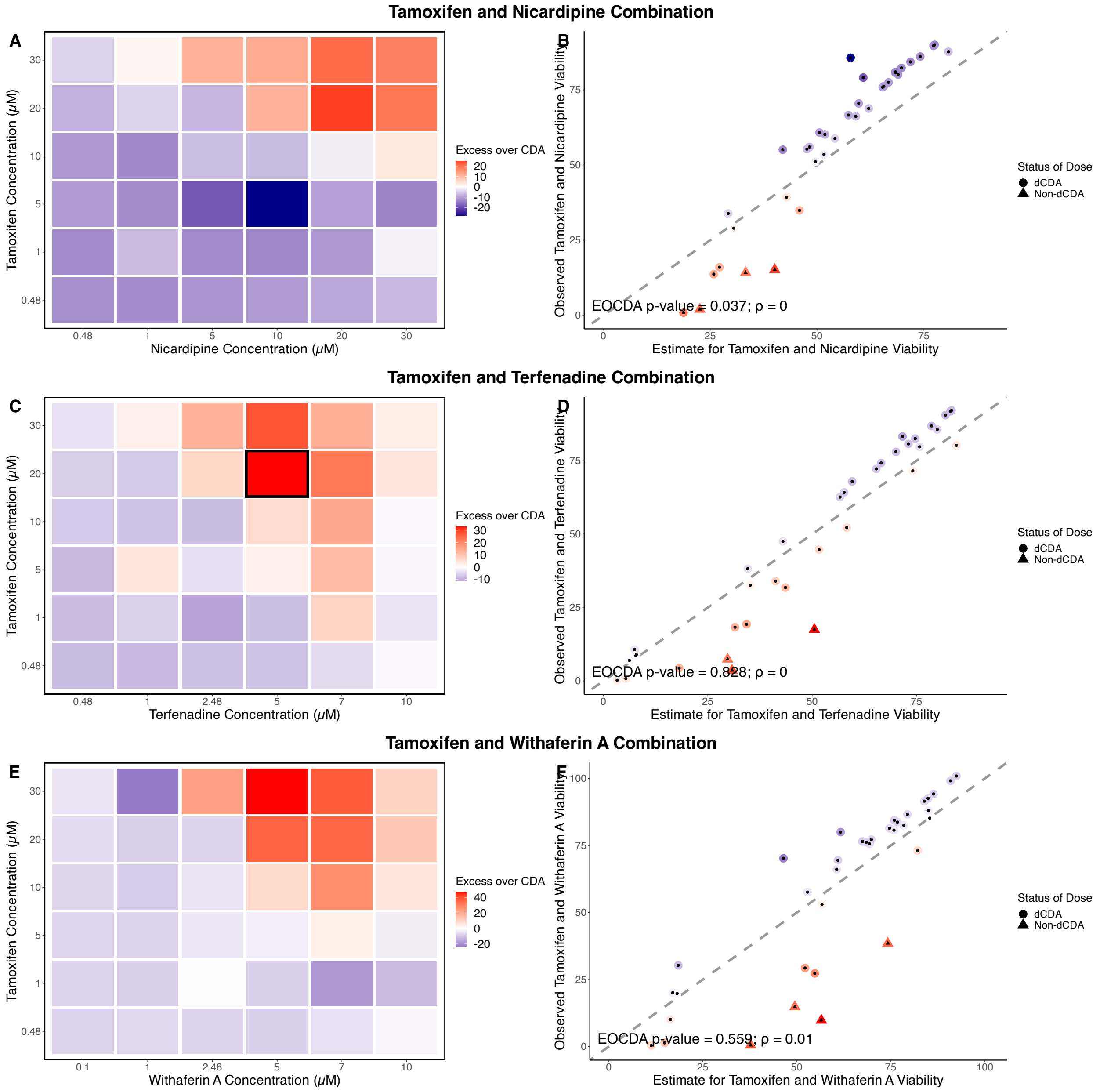


**Figure S12: dCDA model results.** Each row corresponds to a given combination. **A, B)** Tamoxifen and Nicardipine in MCF7 cells. **C, D)** Tamoxifen and Terfenadine combination in MCF7 cells collected after 24 hours. **E, F)** Tamoxifen and Withaferin A in MCF7 cells collected after 24 hours. **A, C, E)** Heatmap of excess over CDA is shown with outlier cells bordered in black. **B, D, F)** Comparison of combination estimates and observed viabilities along with goodness-of-fit (GoF) p-value and corresponding optimal Spearman correlation's estimate. Points are colored with the same scale as its corresponding EOCDA matrix. If the GoF p-value $> 0.01$ for the overall combination, then each point (i.e., dose) is classified as following the dCDA model or not (i.e., likely synergistic or antagonistic behavior) (See Supplement).

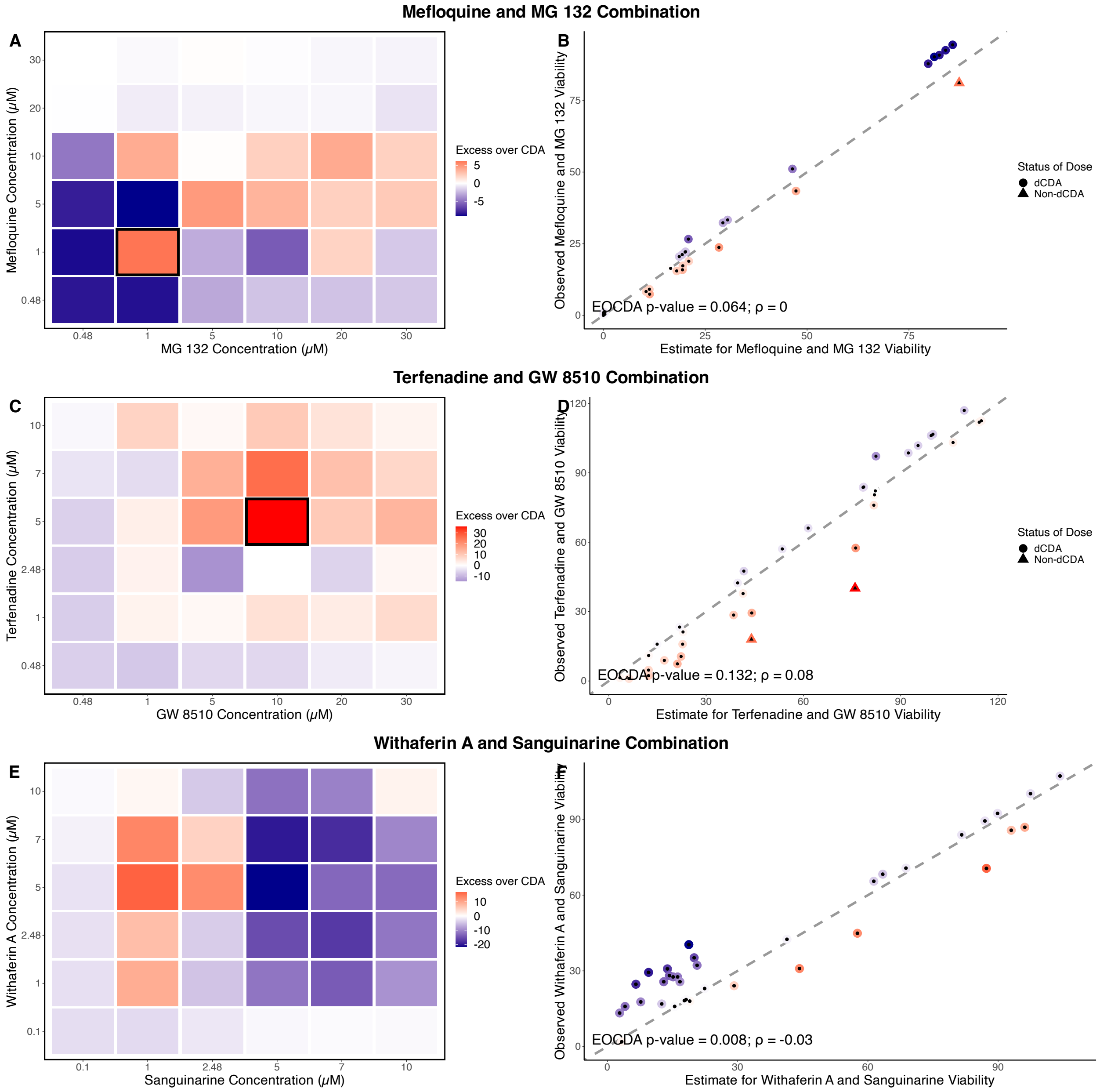


**Figure S13: dCDA model results.** Each row corresponds to a given combination. **A, B)** Tamoxifen and MG 132 in MCF7 cells. **C, D)** GW 8510 and Terfenadine combination in MCF7 cells collected after 24 hours. **E, F)** Sanguinarine and Withaferin A in MCF7 cells collected after 24 hours. **A, C, E)** Heatmap of excess over CDA is shown with outlier cells bordered in black. **B, D, F)** Comparison of combination estimates and observed viabilities along with goodness-of-fit (GoF) p-value and corresponding optimal Spearman correlation's estimate. Points are colored with the same scale as its corresponding EOCDA matrix. If the GoF p-value $> 0.01$ for the overall combination, then each point (i.e., dose) is classified as following the dCDA model or not (i.e., likely synergistic or antagonistic behavior) (See Supplement).

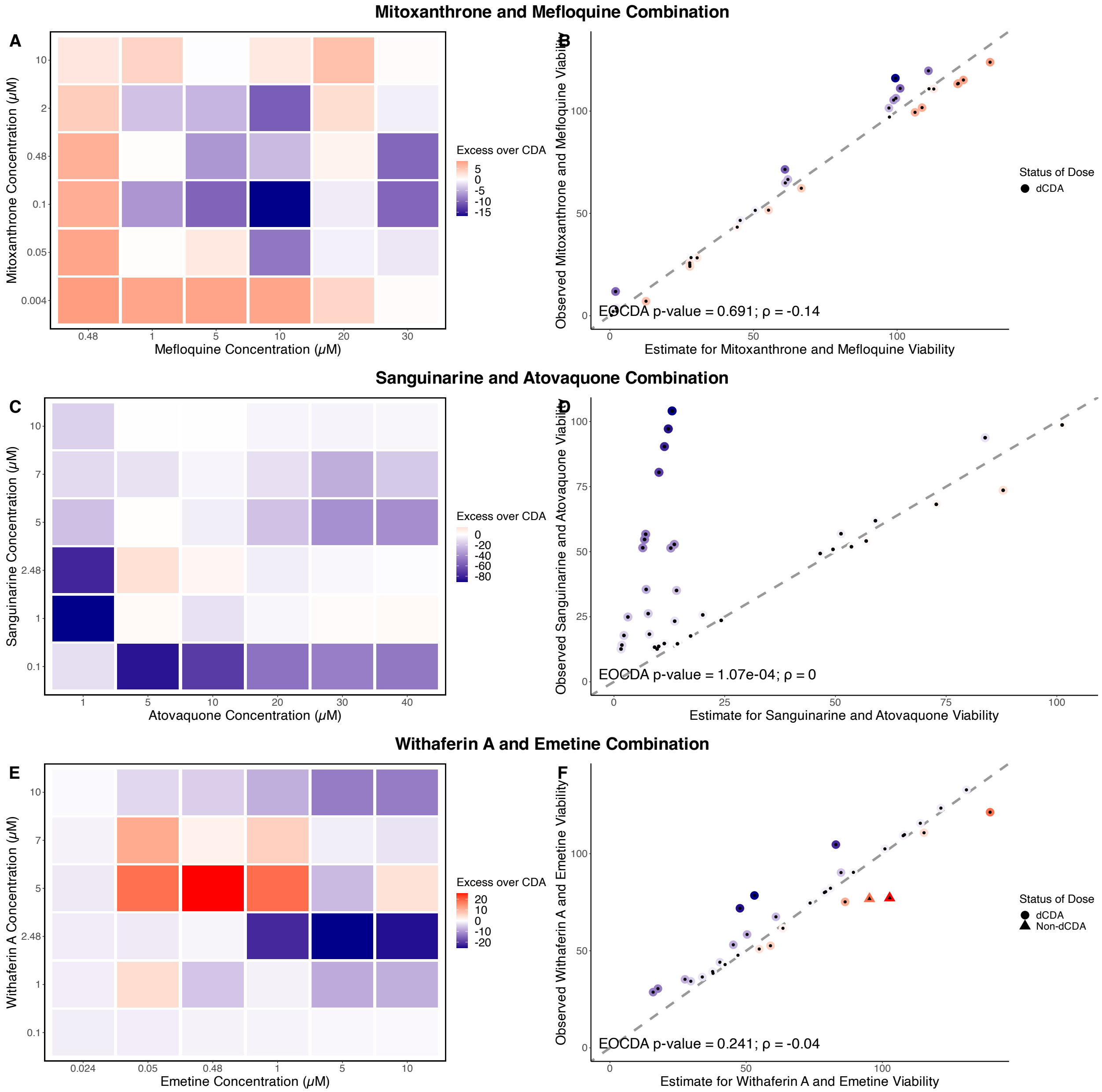


**Figure S14: dCDA model results.** Each row corresponds to a given combination. **A, B)** Mitoxanthrone and Mefloquine in MCF7 cells. **C, D)** Sanguinarine and Atovaquone combination in MCF7 cells collected after 24 hours. **E, F)** Emetine and Withaferin A in MCF7 cells collected after 24 hours. **A, C, E)** Heatmap of excess over CDA is shown with outlier cells bordered in black. **B, D, F)** Comparison of combination estimates and observed viabilities along with goodness-of-fit (GoF) p-value and corresponding optimal Spearman correlation's estimate. Points are colored with the same scale as its corresponding EOCDA matrix. If the GoF p-value $> 0.01$ for the overall combination, then each point (i.e., dose) is classified as following the dCDA model or not (i.e., likely synergistic or antagonistic behavior) (See Supplement).

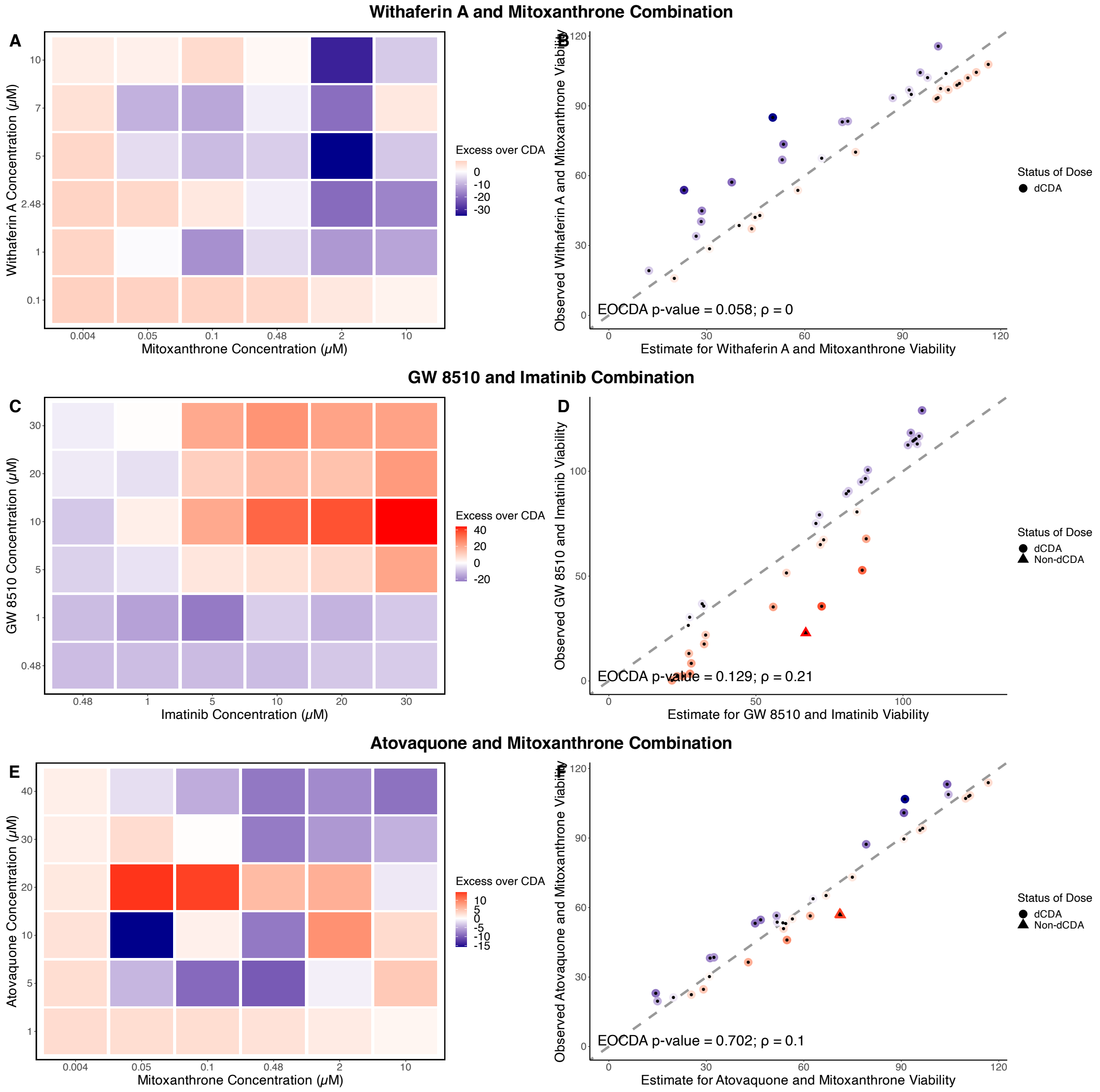


**Figure S15: dCDA model results.** Each row corresponds to a given combination. **A, B)** Withaferin A and Mitoxanthrone combination in MCF7 cells collected after 24 hours. **C, D)** GW 8510 and Imatinib combination in MCF7 cells collected after 24 hours. **E, F)** Atovaquone and Mitoxanthrone combination in MCF7 cells collected after 24 hours. **A, C, E)** Heatmap of excess over CDA is shown with outlier cells bordered in black. **B, D, F)** Comparison of combination estimates and observed viabilities along with goodness-of-fit (GoF) p-value and corresponding optimal Spearman correlation's estimate. Points are colored with the same scale as its corresponding EOCDA matrix. If the GoF p-value $> 0.01$ for the overall combination, then each point (i.e., dose) is classified as following the dCDA model or not (i.e., likely synergistic or antagonistic behavior) (See Supplement).

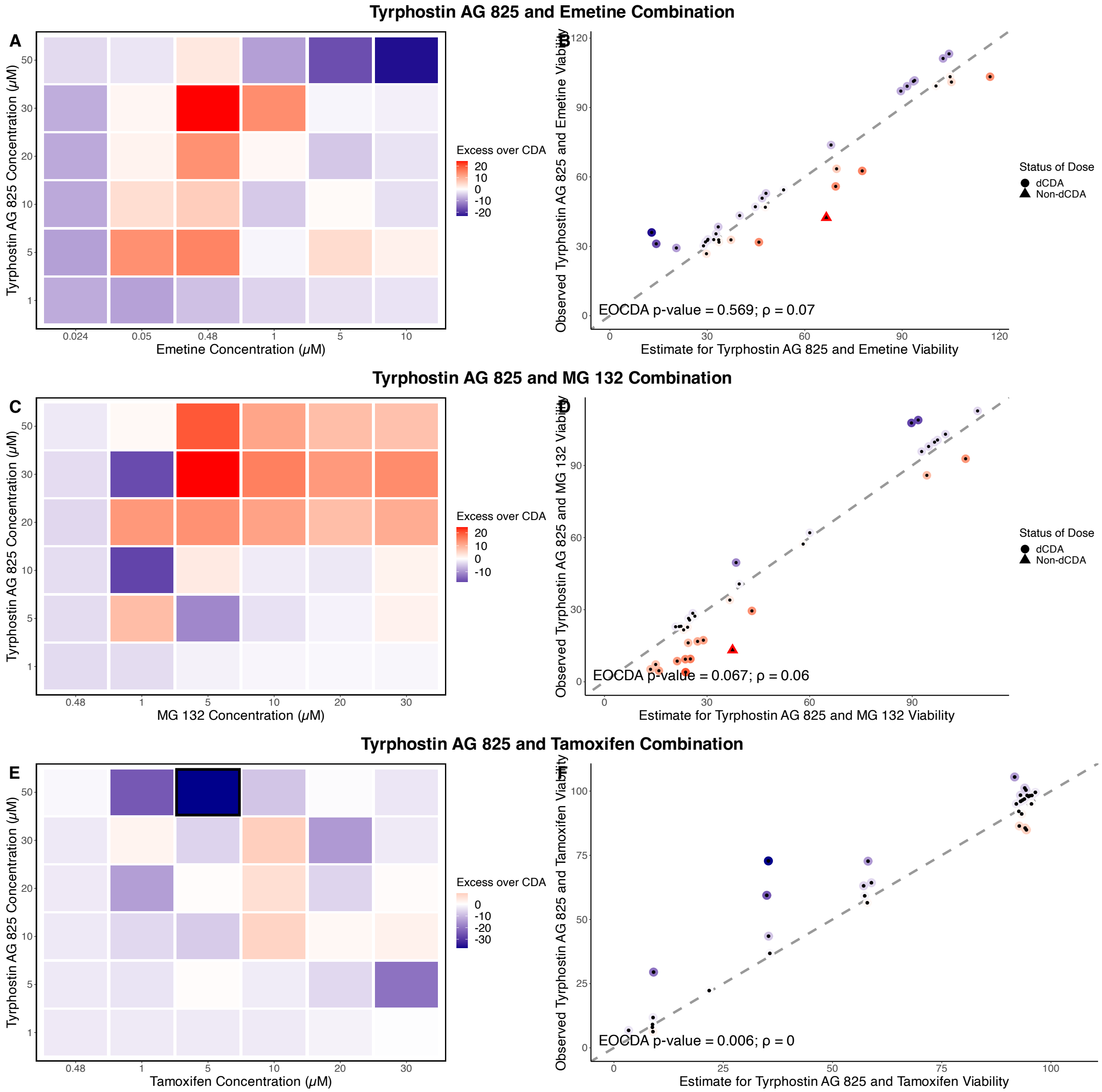


**Figure S16: dCDA model results.** Each row corresponds to a given combination. **A, B)** Tyrphostin AG 825 and Emetine combination in MCF7 cells collected after 24 hours. **C, D)** Tyrphostin AG 825 and MG 132 in MCF7 cells collected after 24 hours. **E, F)** Tyrphostin AG 825 and Tamoxifen combination in MCF7 cells collected after 24 hours. **A, C, E)** Heatmap of excess over CDA is shown with outlier cells bordered in black. **B, D, F)** Comparison of combination estimates and observed viabilities along with goodness-of-fit (GoF) p-value and corresponding optimal Spearman correlation's estimate. Points are colored with the same scale as its corresponding EOCDA matrix. If the GoF p-value $> 0.01$ for the overall combination, then each point (i.e., dose) is classified as following the dCDA model or not (i.e., likely synergistic or antagonistic behavior) (See Supplement).

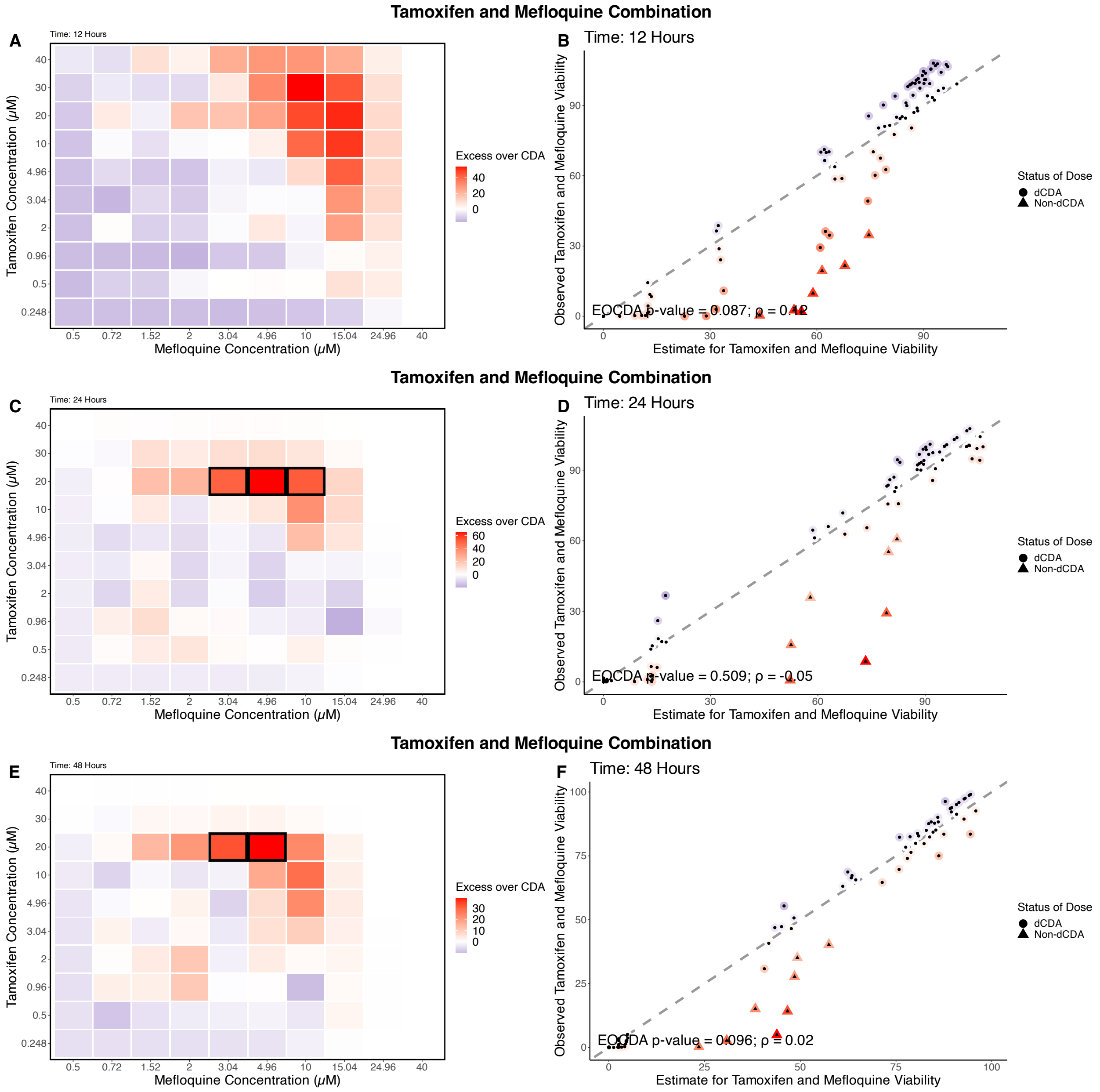


**Figure S17: dCDA model results.** Each row corresponds to a given combination. **A, B)** Tamoxifen and Mefloquine combination in MCF7 cells collected at 12 hours. **C, D)** Tamoxifen and Mefloquine in MCF7 cells collected at 24 hours. **E, F)** Tamoxifen and Mefloquine in MCF7 cells collected at 48 hours. **A, C, E)** Heatmap of excess over CDA is shown with outlier cells bordered in black. **B, D, F)** Comparison of combination estimates and observed viabilities along with goodness-of-fit (GoF) p-value and corresponding optimal Spearman correlation's estimate. Points are colored with the same scale as its corresponding EOCDA matrix. If the GoF p-value $> 0.01$ for the overall combination, then each point (i.e., dose) is classified as following the dCDA model or not (i.e., likely synergistic or antagonistic behavior) (See Supplement).

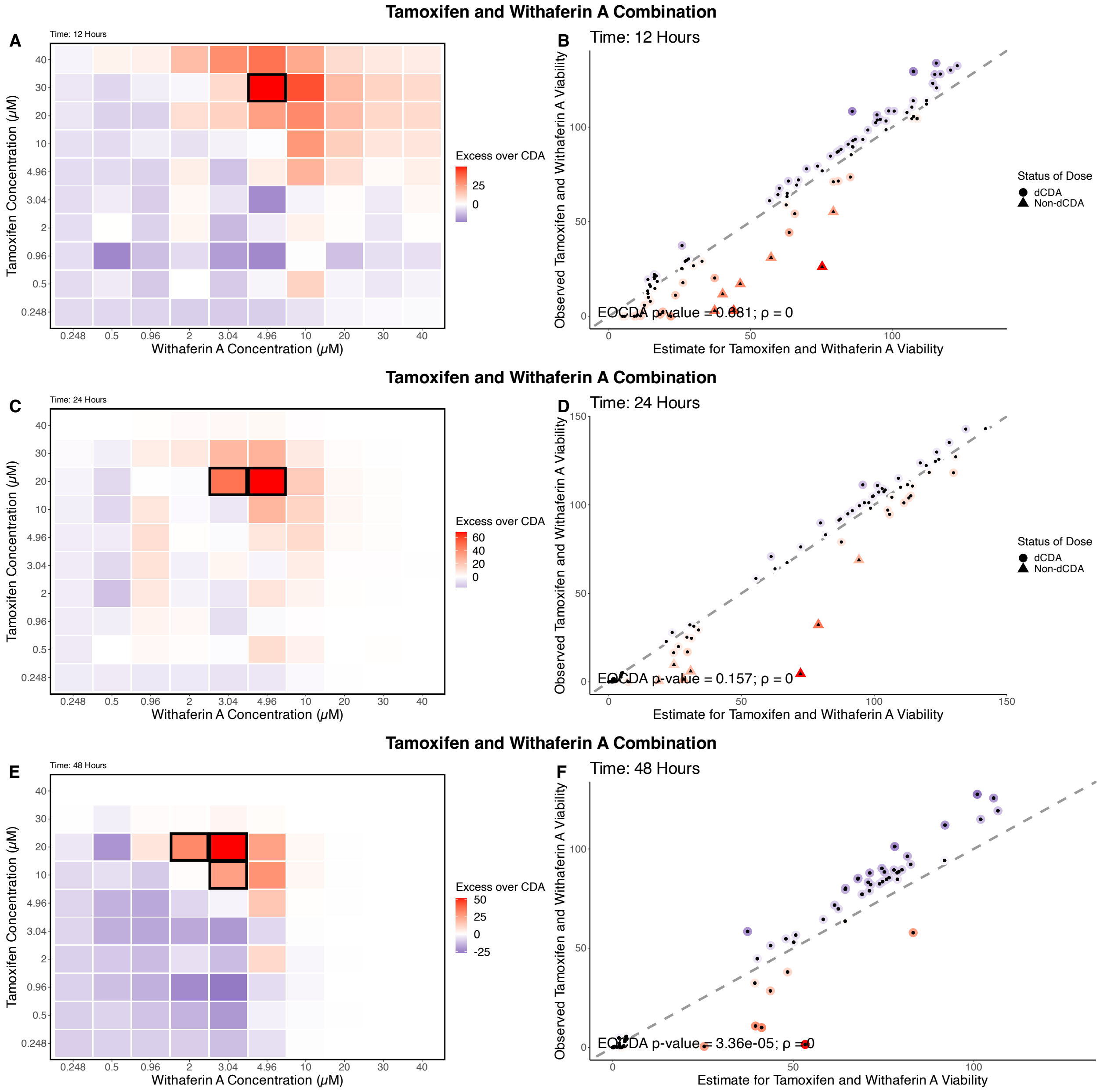


**Figure S18: dCDA model results.** Each row corresponds to a given combination. **A, B)** Tamoxifen and Withaferin A combination in MCF7 cells collected at 12 hours. **C, D)** Tamoxifen and Withaferin A in MCF7 cells collected at 24 hours. **E, F)** Tamoxifen and Withaferin A in MCF7 cells collected at 48 hours. **A, C, E)** Heatmap of excess over CDA is shown with outlier cells bordered in black. **B, D, F)** Comparison of combination estimates and observed viabilities along with goodness-of-fit (GoF) p-value and corresponding optimal Spearman correlation's estimate. Points are colored with the same scale as its corresponding EOCDA matrix. If the GoF p-value $> 0.01$ for the overall combination, then each point (i.e., dose) is classified as following the dCDA model or not (i.e., likely synergistic or antagonistic behavior) (See Supplement).
